# Bilingual language processing relies on shared semantic representations that are modulated by each language

**DOI:** 10.1101/2024.06.24.600505

**Authors:** Catherine Chen, Xue L. Gong, Christine Tseng, Daniel L. Klein, Jack L. Gallant, Fatma Deniz

**Affiliations:** Electrical Engineering and Computer Science, University of California, Berkeley, CA 94720, USA; Helen Wills Neuroscience Institute, University of California, Berkeley, CA 94720, USA; Faculty of Electrical Engineering and Computer Science, Technische Universität Berlin, 10623 Berlin, Germany; Bernstein Center for Computational Neuroscience, 10115 Berlin, Germany

## Abstract

Billions of people throughout the world are bilingual, and they can extract meaning from multiple languages. While some evidence suggests that there is a shared system in the human brain for processing semantic information from native and non-native languages, other evidence suggests that semantic processing is language-specific. We conducted a study to determine how semantic information for different languages is represented in the brains of bilinguals. Functional magnetic resonance imaging (fMRI) was used to record brain responses while participants read several hours of natural narratives in their native (Chinese) and non-native (English) languages. These data were then used to compare semantic representations between the two languages. We find that semantic representations are largely shared between languages, but that there are fine-grained differences in the representation of some semantic categories across languages. These results reconcile current competing theories of bilingual language processing.

**Significance Statement:** Bilinguals understand the meaning of words in multiple languages. Whether this capacity reflects a shared brain system for processing both native and non-native languages, or whether processing is language-specific is still unclear. Here, we examine whether and how semantic representations in the brain support shared and/or language-specific processing. We recorded brain activity from participants reading narratives in their native (Chinese) and non-native (English) languages, and modeled how their brains represent word meaning in each language. We show that semantic representations are similar between the two language conditions, and that these representations are systematically modulated between native and non-native language comprehension.

## Introduction

Over four billion people throughout the world are bilingual, and they can understand the meaning of words in both their native and non-native languages (1). However, there is conflicting evidence about how bilinguals process the meanings of words from different languages. Some behavioral studies have shown that knowledge of one language can interfere with semantic processing in another. For example, second language acquisition can increase processing times for false cognates (2–4), and it can change how speakers describe concepts in their native language (5). This interference suggests that semantic processing is shared between languages.

Other behavioral studies have shown that semantic processing differs between languages. For example, native and non-native languages differ in perceived emotional intensity (6, 7), memory storage and recall during conversation (8), and the speed of numerical processing (9, 10). From behavioral evidence alone, it remains unclear how semantic representations in the brains of bilinguals support both shared and language-specific processing.

Several prior neuroimaging studies of bilingualism have argued that different languages are represented in a common set of brain regions (11–17), though a few studies have reported that different languages are represented in anatomically distinct regions (11–13). However, these prior studies have not used methods that were sensitive enough to truly resolve this issue. Many of the prior studies used contrast-based methods that can identify which rain regions are activated during language comprehension (11–17), but which cannot reveal precisely how semantic information is represented therein. This is a critical problem, because semantic information in language is represented in intricate cortical maps, and so different word meanings elicit different patterns of brain activity across the cerebral cortex (18, 19). Thus, determining whether semantic representations are similar or different between languages requires not only identifying regions of activation, but also mapping functional semantic representations in each language. Furthermore most prior functional imaging studies of bilingualism did not perform within-participant comparisons between languages (11, 20–22), which made it impossible to overcome individual differences in brain function and anatomy (23) in order to recover fine-grained functional maps of semantic representation. Finally, many prior studies used controlled stimuli (11, 15–17, 24) that do not elicit the same type of robust, widespread activity as occurs during natural language comprehension (25, 26). Thus, it is unclear whether prior results generalize to naturalistic settings (25).

A new hypothesis for how semantic representations in the brains of bilinguals might support both shared and language-specific processing comes from (26, 27). It is thought that these tuning shifts optimize the representation of task-relevant semantic information, at the cost of irrelevant information. Importantly, even when semantic tuning remains largely consistent between task conditions, small, measurable tuning shifts often still occur. Based on these prior studies we hypothesized that native and non-native languages themselves act as different task contexts that will modulate semantic representations in the brain. Under this hypothesis, even if semantic representations are largely similar between native and non-native languages, there could still be significant shifts in semantic tuning across languages that reflect language-specific representations of word meaning (Figure 1A).

**Figure 1.**
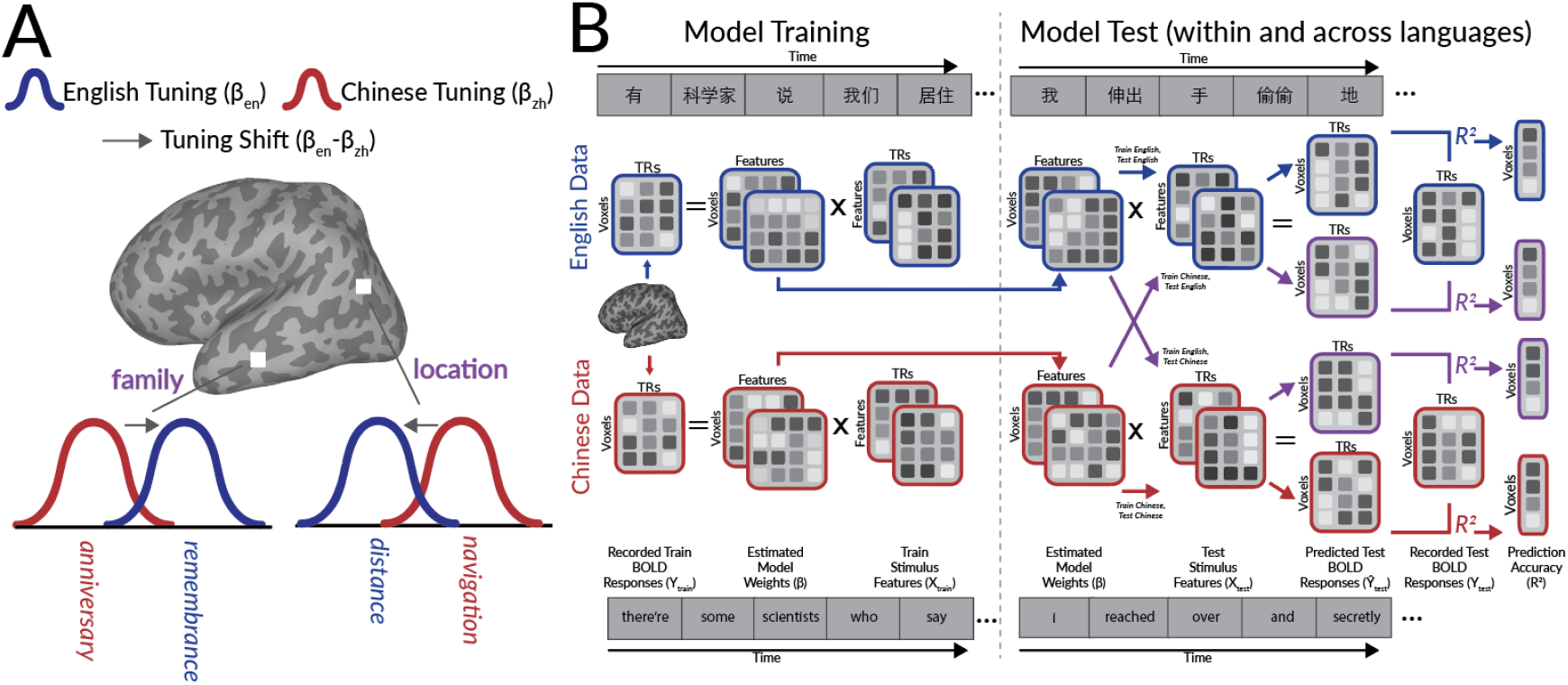
Hypothesis and experimental procedure. **A**. Schematic illustrating hypothesized semantic representation shifts. The semantic tuning of a voxel describes how it responds to word meaning. Shifts in semantic representations were quantified as the change in semantic tuning between languages. For each voxel, blue and red curves respectively denote semantic tuning in L1 (Chinese) and L2 (English). Arrows represent semantic tuning shifts from Chinese to English. The hypothetical voxel in the parietal cortex responds to location-related words in both languages, but emphasizes number-related aspects (“distance”) in English, and action-related aspects (“navigation”) in Chinese. The hypothetical voxel in temporal cortex responds to family-related words in both languages, but emphasizes emotion-related aspects (“remembrance”) in English, and number-related aspects (“anniversary”) in Chinese. **B**. Experiment and modeling procedure. Six fluent Chinese-English bilingual participants read over two hours of narratives in each language while BOLD responses were measured using fMRI. Semantic stimulus features were constructed by projecting each stimulus word into a 300-dimensional word embedding space (28). Ridge regression was used to estimate encoding models that describe voxelwise semantic tuning for each participant and language. Estimated model weights were used to predict BOLD responses to held-out narratives not used for model estimation. Model weights estimated in one language were used to predict BOLD responses to the same language (red and blue arrows; *within-language*) and to the other language (purple arrows; *across-language*). Prediction accuracy was quantified as the coefficient of determination ( *R*^2^) between predicted and recorded BOLD responses.

To test this hypothesis, we designed a study to investigate how the brains of bilinguals encode the meanings of words in their native and non-native languages. Six participants who are fluent in both Chinese (native; L1) and English (non-native; L2) read natural narratives for over two hours in each language while functional magnetic resonance imaging (fMRI) was used to record brain activity. Voxelwise encoding models were used to recover semantic representations for each language separately. Semantic representations were compared between the two languages for each individual participant separately. Our results support the hypothesis that semantic brain representations are largely shared between languages, but that these shared representations are modulated by each language.

## Results

Six Chinese-English bilinguals read natural narratives while fMRI was used to record blood oxygen level dependent (BOLD) responses. Each narrative was presented in both English and Chinese as written text. Words were presented one at a time at a natural reading rate. Each participant read narratives for over two hours per language. Semantic stimulus features were extracted by projecting each word into a distributional word embedding space (fastText (28, 29)) in which words in different languages with similar meanings project to nearby vectors. Regularized regression was used to estimate voxelwise encoding models that describe how the semantic stimulus features are encoded in brain responses. Models were estimated separately for each participant and language (Figure 1B). Nuisance features such as word rate and spatiotemporal visual features were regressed out of BOLD responses prior to model estimation. The estimated model weights describe the semantic tuning of each individual voxel, for each language separately. Estimated model weights were compared between English and Chinese to examine whether and how semantic representations differ between languages. To ensure the generalizability of the results and to prevent overfitting, the estimated model weights were evaluated on BOLD responses to a held-out story that was not used for model estimation. To ensure generalization to new participants, models were estimated and evaluated separately for each participant and the results were replicated in each individual participant, and data for two participants (P5 and P6) were not analyzed until the entire analysis pipeline was finalized.

The hypothesis that semantic representations are similar between languages presents two predictions. First, the same brain regions should represent semantic information for both languages. Second, within these regions the same word meaning should activate a similar set of voxels across languages. To study the first prediction, we examined the prediction accuracy of the estimated semantic model weights. If a voxel represents semantic information in one language, then estimated semantic model weights for that language will accurately predict held-out test data for that language. Prediction accuracy was computed as the coefficient of determination ( *R*^2^) between predicted and recorded BOLD responses for each voxel, participant, and language separately.

Group-level results were computed by projecting voxelwise accuracies for each participant to a template space (fsAverage (30)), and then averaging the projected values across participants for each vertex and language separately. Figure 2A shows vertexwise group-level prediction accuracy for each language separately. Results for each participant are similar to group-level results (Figure S5 and S6). For each language, prediction accuracy is highest within bilateral temporal, parietal, and prefrontal cortices, regions that together comprise the *semantic system* (18, 31). Figure 2B directly compares the group-level prediction accuracy of each vertex between the two languages. Prediction accuracy was significantly correlated between the two languages both at the group-level (r=.68, one-sided p<.001 by a permutation test) and for each individual participant (r=.49, .36, .28, .32, .19, .46 for P1-P6 respectively, for each participant p<.001 by a one-sided permutation test). These results show that the same brain regions represent semantic information for both languages.

**Figure 2.**
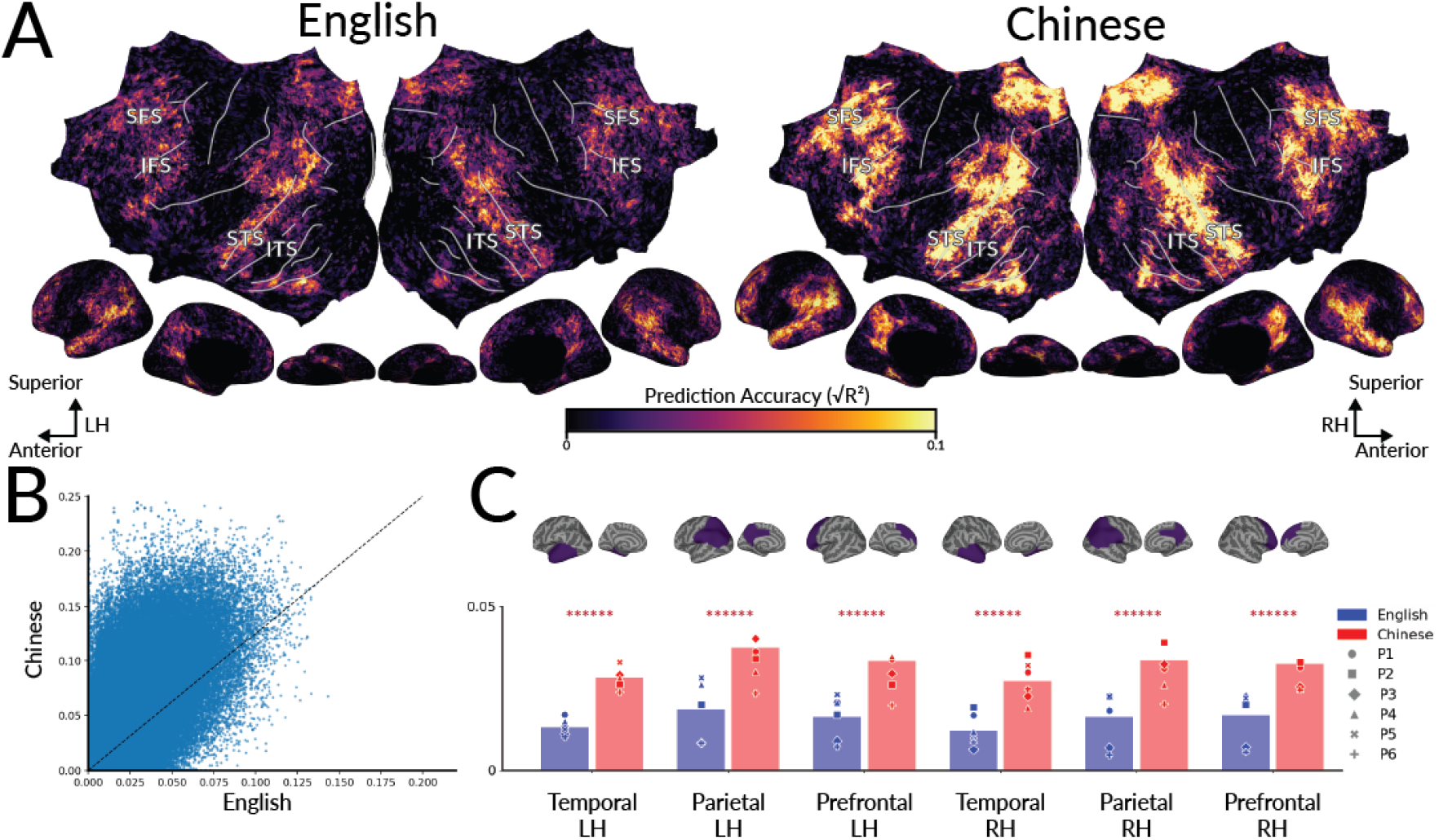
Cortical distribution of semantic representations for each language. To determine where semantic information is represented for each language, voxelwise models estimated for each language were used to predict held-out data for the same language. Prediction accuracy was computed as the coefficient of determination (*R*^2^) between predicted and recorded BOLD responses. **A**. Group-level prediction accuracy. Results are shown for each language on the flattened cortical surface of the template space. For both languages, prediction accuracy is highest in the bilateral temporal, parietal, and prefrontal cortices. Prediction accuracy in these regions is statistically significant for each participant (Figure S6). The same brain regions are well-predicted for both languages. (SFS=superior frontal sulcus; IFS=inferior frontal sulcus; STS=superior temporal sulcus; ITS=inferior temporal sulcus, LH=left hemisphere; RH=right hemisphere) **B**. Group-level prediction accuracy for English (x-axis) vs Chinese (y-axis). Each point represents one vertex. Prediction accuracy is significantly positively correlated between the two languages (r=.68, one-sided p < .001 by a permutation test). **C**. Prediction accuracy by cortical region. For each brain region and participant, blue and red markers show the mean prediction accuracy over voxels for English and Chinese respectively. Bars show the mean across participants. Red asterisks denote the number of participants for which prediction accuracy is significantly higher in Chinese than in English (one-sided p<.05 by a permutation test). While the same brain regions are well-predicted for both languages, the semantic model explains more variability in brain responses for Chinese (L1) than for English (L2).

While the same brain regions are well-predicted for both languages, visual inspection of Figure 2A indicates that prediction accuracy in these participants is overall higher in Chinese than in English. To quantify this difference for each brain region in the semantic system, we identified the set of voxels in the temporal, parietal, and prefrontal regions for each participant using the Desikan-Killiany atlas (32). We then computed the average prediction accuracy separately for each language and region. Figure 2C shows the average prediction accuracy for each language, region, and participant. Prediction accuracy within these regions is significantly greater in Chinese than in English (one-sided p<.05 by a permutation test for all brain regions and participants).

There are two potential explanations for why prediction accuracy is higher in Chinese (L1) than in English (L2) in this experiment. One possibility is that the recorded brain responses simply have a higher signal-to-noise ratio (SNR) in Chinese than in English. This could happen if, for example, the participants paid more attention when the stories were presented in Chinese than in English. To test this possibility, we examined voxelwise explainable variance (EV). EV estimates the SNR of each voxel by measuring the repeatable signal over multiple repetitions of the same stimulus (33–35). If the SNR is higher for Chinese than for English, then EV will be higher in Chinese than in English. Across participants, EV was not higher in Chinese than in English (Figure S7), suggesting that SNR is not higher in Chinese than in English.

A second possibility is that SNR is comparable between languages, but our semantic model better represents brain responses for Chinese as compared to English. Brain representations reflect not only semantic processes, but also other processes such as sentence-level syntactic parsing, working memory load, or high-level control. The semantic model weights can only predict how the brain represents information captured by the word embeddings, and the word embeddings do not capture extra-linguistic processes such as working memory load. Aspects of language processing that are not reflected in the word embeddings contribute to variance in the repeatable signal that is not explained by the semantic model weights. To test for this possibility, we examined voxelwise noise-ceiling corrected prediction accuracy. Noise-ceiling corrected prediction accuracy measures how well a model predicts brain responses, relative to the maximum possible accuracy of a model given the noise in the data (33–35). If the semantic model better represents brain responses for Chinese, then the noise-ceiling corrected prediction accuracy will be higher for Chinese than for English. Across participants, the noise-ceiling corrected prediction accuracy was significantly higher for Chinese than for English (Figure S8). Thus, the semantic model predicts a higher fraction of explainable signal in Chinese than in English. This could indicate that semantic processes more strongly influence brain representations for Chinese (participants’ L1) as compared to English (L2).

Differences in noise-ceiling corrected prediction accuracy could be a result of differences between native and non-native language processing, or because of differences that are intrinsic to Chinese and English language processing. Future work with broader bilingual populations could help distinguish between these two possibilities.

Figure 2 shows that between languages semantic information is represented in the same brain regions. The hypothesis that semantic representations are similar between languages additionally suggests that within these brain regions, voxels should be tuned towards similar word meanings in different languages. Alternatively, voxels could instead be tuned towards different word meanings between different languages. For example, a voxel might be tuned towards emotion-related words in one language and location-related words in the other. Although such a voxel would represent semantic information in both languages, its semantic tuning would differ between languages. To examine whether the semantic tuning of each voxel is similar between languages, we classified voxels into semantic clusters based on semantic tuning in each language, and then evaluated whether voxelwise cluster assignments are similar between languages.

First, we used a cross-validated clustering approach (*model connectivity (36)*) to identify semantic clusters from the estimated semantic model weights. Five clusters best summarized the distribution of semantic model weights across participants and languages (Figure S9). To interpret each cluster, we identified the English stimulus words that are closest to each cluster. Distance between a word and a cluster was computed as the Pearson correlation between the word embedding and the cluster centroid. Figure 3A lists the closest words to each cluster. The clusters categorize voxels that are tuned towards words related to family (Cluster 1), communication (Cluster 2), cognition (Cluster 3), locations (Cluster 4), and numbers/names (Cluster 5). Then each voxel was assigned to the semantic cluster with the lowest Euclidean distance between the cluster centroid and the vector of semantic model weights for that voxel. Cluster assignments were performed for each participant and language separately. To summarize results across participants, we projected semantic model weights of each participant and language to the template space. Then for each vertex of the template space and for each language separately, we computed the mean model weights across participants, and used the mean model weights to assign each vertex to one of the five clusters.

**Figure 3.**
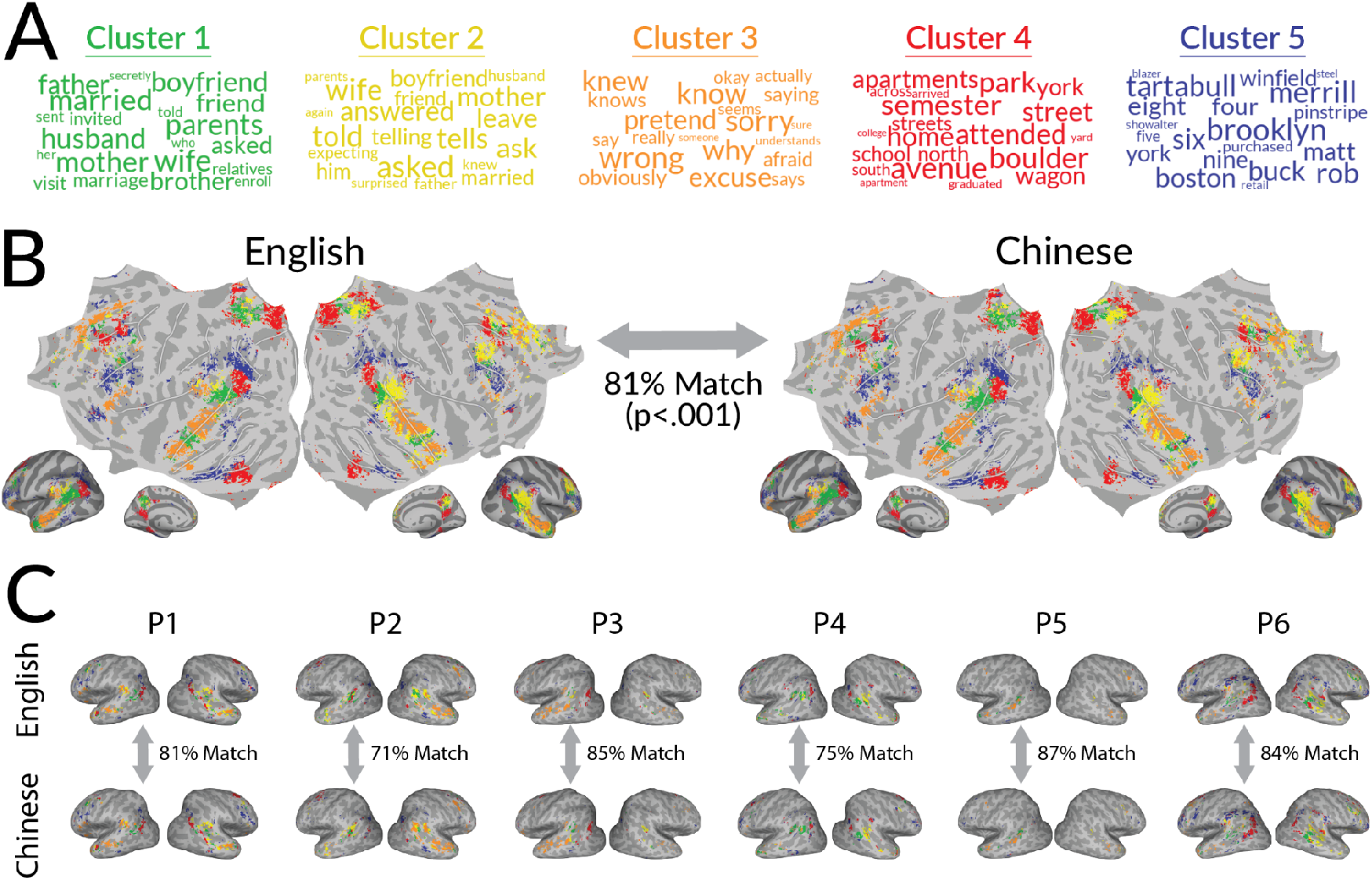
Shared semantic representations between languages. Hierarchical clustering on estimated semantic model weights was used to categorize voxels based on their semantic tuning in each language. **A**. Words closest to each cluster. The five semantic clusters are respectively related to family (Cluster 1, green), communication (Cluster 2, yellow), cognition (Cluster 3, orange), locations (Cluster 4, red), and numbers/names (Cluster 5, blue). **B**. Cortical distribution of semantic clusters across all participants. Group-level results are shown for each language. Vertex color reflects the assigned semantic cluster. Vertices that are poorly-predicted by the semantic model are shown in grey. Cluster assignments are the same between languages in 81% of well-predicted vertices. **C**. Cortical distribution of semantic clusters in individual participants. Voxel color reflects the assigned semantic cluster, following the same color scheme as in B. For each participant, cluster assignments are the same between languages in over 70% of well-predicted voxels. Thus, semantic representations are largely shared between languages in individual participant and group-level.

Figure 3B shows semantic cluster assignments for each language at the group-level. Semantic cluster assignments are shown for vertices that were well-predicted in both languages (√*R*^2^ > 0. 1) in at least one participant. To quantify whether the cluster assignments are consistent between languages, we computed the confusion matrix between cluster assignments in English and Chinese. For 81% of well-predicted vertices semantic cluster assignments are the same between languages (p<.001 by a permutation test). Figure 3C shows semantic cluster assignments for each language in individual subjects. For each participant, cluster assignments are the same between languages for over 70% of well-predicted voxels (p<.001 by a permutation test). The consistency of semantic cluster assignments suggests that semantic tuning is similar between languages. As a converging test, we measured whether model weights estimated in one language could predict voxel responses to the other language (across-language prediction accuracy). Models estimated for English accurately predicted brain responses to Chinese and vice versa throughout bilateral temporal, parietal, and prefrontal cortices (Figures S6, S10). Overall, these results show that semantic representations are similar between languages.

Based on prior behavioral evidence that semantic processing is to some extent different between languages (6–8) we hypothesized that semantic representations in the brain are modulated by each language (Figure 1A). Modulation of semantic representations could be observed as a shift in the exact semantic tuning of each voxel. For example, a voxel might respond to the same semantic clusters (location-related words) in both languages (Figure 2 and 3), but the semantic tuning of the voxel may subtly shift to respond more strongly to words associated with actions (“navigation”) in one language, and words associated with numbers (“distance”) in the other.

To determine whether the semantic tuning of each voxel shifts between languages, we investigated the change in estimated model weights between languages 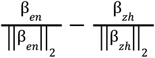, which we refer to as the voxelwise *semantic tuning shift*. The semantic tuning shift for each voxel describes which word meanings are emphasized in each language relative to the other language. To identify dimensions of semantic tuning shifts that are reliable across voxels and participants, we used a leave-one-participant-out procedure. For each of the six participants, we concatenated voxelwise semantic tuning shifts from the other five participants and then used principal component analysis (PCA) to obtain 300 orthogonal axes (principal components; PCs). Then we evaluated how well each PC explains variance in semantic tuning shifts for the left-out participant. The chance rate was defined as the variance explained by the PCs of semantic model weights within each language. The first semantic tuning shift PC, which is the main dimension along which semantic representations shift between languages, reliably explains more variance than chance (Figure S11, by a 95% bootstrap confidence interval). To obtain an estimate of the first semantic tuning shift PC that incorporates data from all six participants, we applied PCA to voxelwise semantic tuning shifts concatenated over all six participants.

The first semantic tuning shift PC reveals the primary dimension in the word embedding space along which voxelwise semantic tuning tends to shift. However, dimensions of the word embedding space are not constructed in terms of classical semantic dimensions. Therefore to understand what a shift toward either end of the PC means in terms of word meanings, we inspected the set of words that are closest to each end of the PC. These words were identified by finding the word embeddings with the highest Pearson correlation with each end of the PC. Figure 4A shows the English stimulus words that are closest to each end of the PC. (The Chinese stimulus words closest to each end of the PC are similar and are shown in Figure S13). The words closest to one end of the PC (colored in purple) are related to numbers and collections (e.g., “three”, “both”). The words closest to the other end (colored in green) are related to actions and human relationships (e.g., “leave”, “boyfriend”). A shift along the PC may reflect higher sensitivity to the meanings of words from certain semantic categories as well as to the meanings of words from certain grammatical categories. For example, between the two ends of the PC one end (colored in purple) includes relatively more quantifiers, while the other end (colored in green) includes relatively more verbs, suggesting that semantic tuning shifts may be tied to the meanings of words in different grammatical categories. For brevity, we describe a semantic tuning shift towards the one end of the PC as an emphasis on number and collection-related word meanings, and shifts towards the other end as an emphasis on action- and relationship-related word meanings. The labels for each end of the PC provide an intuitive description of how the meaning of words in different languages can shift the representation of a voxel. However, these labels may not fully capture the precise description of the semantic tuning shifts along each axis of the PC. For example, a shift along the PC could reflect not only differences in sensitivity to the meaning of words from intuitive semantic categories, but also other factors such as the sensitivity to the meaning of words from different grammatical categories.

**Figure 4.**
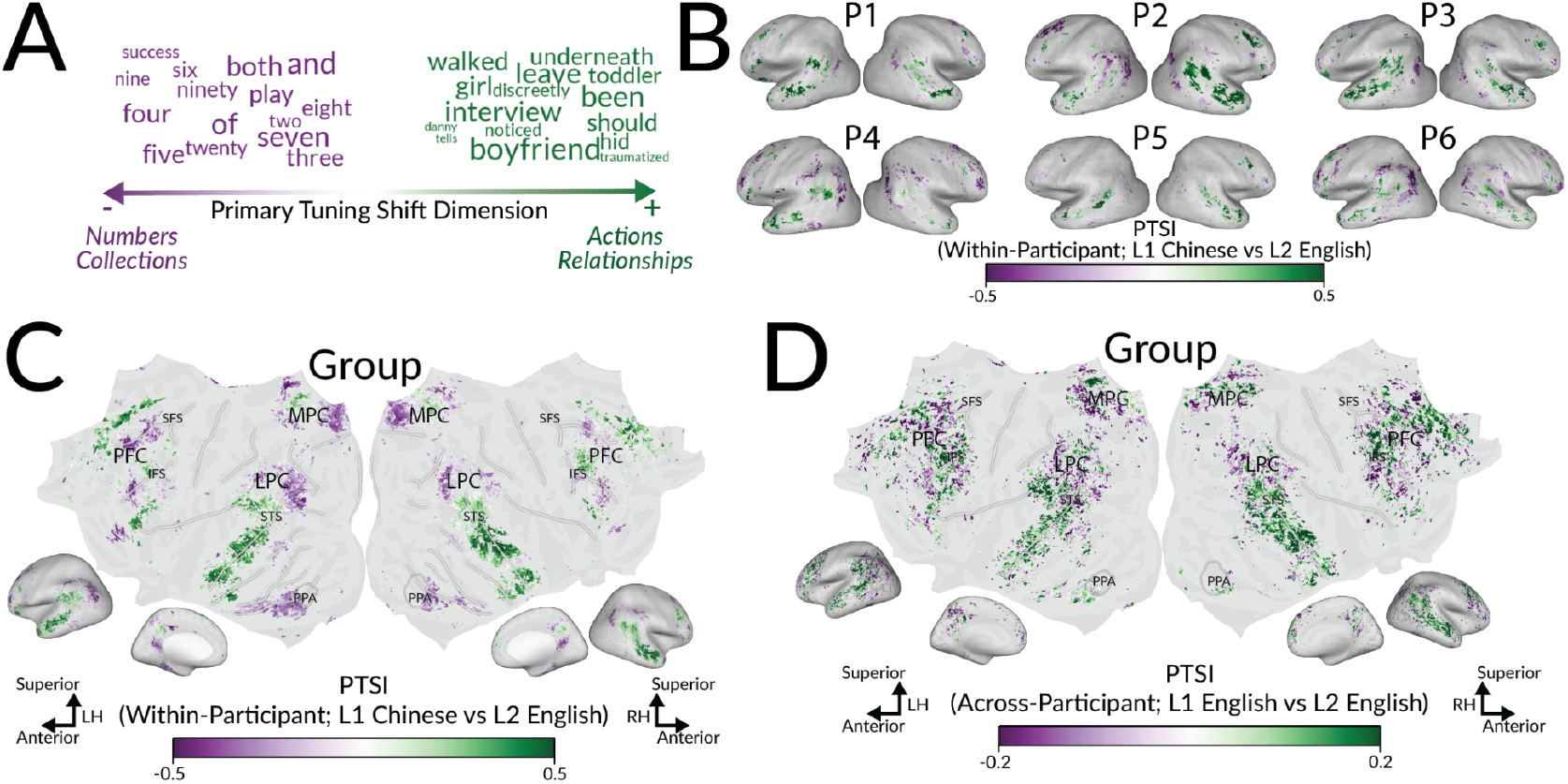
Cortical distribution of semantic tuning shifts between languages. Voxelwise *semantic tuning shift* was defined as the change in model weights from Chinese (L1) to English (L2): 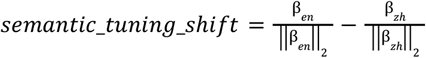. These shifts in semantic tuning alter the voxel’s sensitivity to certain word meanings. The primary dimensions of voxelwise semantic tuning shifts were identified using principal component analysis (PCA). The first semantic tuning shift PC reliably explains variance in semantic tuning shifts across voxels and participants (Figure S11). **A**. Interpretation of the first PC. Words closest to one end of the PC are shown in purple. Words closest to the other end of the PC are shown in green. One end emphasizes number- and collection-related semantics (purple), while the other end emphasizes action- and relationship-related semantics (green). **B**. Cortical distribution of semantic tuning shifts for each participant. The direction of voxelwise semantic tuning shifts was summarized with the *primary tuning shift index* (PTSI), which is the Pearson correlation between a voxel’s semantic tuning shift and the first semantic tuning shift PC. The color of each voxel reflects its PTSI. Voxels shown in purple shift towards the negative end of the PC. Voxels shown in green shift towards the positive end of the PC. Poorly predicted vertices are shown in grey. The cortical distribution of PTSI appears to be consistent across participants. **C**. Cortical distribution of semantic tuning shifts at the group-level. Group-level PTSI is shown on the flattened cortical surface of the template space. The color of each vertex reflects group-level PTSI, according to the same colormap as in B. The cortical distribution of group-level PTSI is consistent with results for each individual participant (p<.05 by a one-sided t-test after Fisher z-transformation, for both positive and negative PTSI all participants except negative PTSI in P5; Figure S12). **D**. Vertexwise semantic tuning shifts between L1 English versus L2 English speakers, with the same English stimuli for all participants. Group-level results are shown on the flattened cortical surface of the template space, according to the same colormap as in C. Vertexwise PTSI between L1 and L2 English speakers is significantly correlated with those shown in C (r=.63; p<.001 by a one-sided permutation test). Overall, these results show that there are systematic shifts in semantic tuning between languages.

To determine whether semantic tuning shifts are consistent across participants, we mapped the direction of semantic tuning shifts onto the cortical surface of each subject. For each voxel we computed the Pearson correlation between the voxel’s semantic tuning shift vector and the first semantic tuning shift PC. We refer to this correlation as the *primary tuning shift index* (PTSI). Because the orientation of PCs are arbitrary, the sign of the PTSI (positive or negative) is also arbitrary. We use the terms “positive” and “negative” simply to distinguish between shifts towards the two ends of the PC. Figure 4B shows voxelwise PTSI for each individual participant. Group-level results are shown in Figure 4C. Voxels with negative PTSI (shown in purple) emphasize number and collection-related words in English, or action and relationship-related words in Chinese.

Negative PTSI is found in bilateral lateral parietal cortex (LPC), fusiform gyrus and near parahippocampal place area (PPA), superior and inferior medial parietal cortex (MPC), and middle frontal cortex (FC). Voxels with positive PTSI (shown in green) emphasize action and relationship-related words in English, or number and collection-related words in Chinese. Positive PTSI is found in bilateral STS, LPC, middle MPC, superior frontal gyrus (SFG), and inferior frontal gyrus (IFG). Visual inspection suggests that the PTSI of each participant is consistent with the rest of the group. To quantify this consistency, we projected voxelwise PTSI of each participant onto the template space, and then compared PTSI between participants in the template space (Figure S12). Vertices with positive PTSI in the rest of the group also have positive PTSI for each individual participant (p<.05, by a one-sample, one-sided t-test after Fisher z-transformation). Vertices with negative PTSI in the rest of the group also have negative PTSI for each individual participant (p<.05, by a one-sample, one-sided t-test after Fisher z-transformation; except P5). The distribution of PTSI is consistent between participants, suggesting that semantic tuning shifts are consistent across individuals. Overall, these results show that there are systematic shifts in semantic tuning between languages and that these shifts are consistent across participants.

There are several potential confounds that could affect the results described in Figure 4. First, the idiosyncrasies of one particular word embedding space (fastText embeddings) could bias the estimated semantic tuning shifts. To examine this possibility, we replicated our analyses with a separate semantic embedding space based on a multilingual language model (mBERT (37)) that differs from fastText in its training objectives, training data, and embedding dimensionality. Semantic models estimated with mBERT produced similar results to those estimated with fastText (Figures S1-S4), suggesting that our results are not confounded by the choice of embedding space.

Second, the estimated semantic tuning shifts could reflect imperfections in word embedding alignments between languages. Because of non-isometry between embedding spaces of different languages and limitations to multilingual alignment procedures, embedding spaces are not perfectly aligned between languages (38).

Thus, it could be that estimated semantic tuning shifts merely reflect imperfect alignments. To examine this possibility, we used English-Chinese translation pairs, and for each pair words computed the difference between the embeddings of the two words. We refer to these differences as *misalignments*. Then we computed the PCs of these misalignments, and examined how well the misalignment PCs explain variance in the estimated semantic tuning shifts. Figure S11 shows that the misalignment PCs do not explain more variance in the estimated semantic tuning shifts than chance (by a 95% bootstrap confidence interval), suggesting that our results are not confounded by imperfections in word embedding alignments.

Third, the estimated semantic tuning shifts could be confounded by encoding model prediction accuracies being higher in Chinese than in English (Figure 2). To examine this possibility, we tested whether semantic tuning shifts were consistent between the subset of voxels where prediction accuracy is higher in Chinese than in English, the subset of voxels where prediction accuracy is higher in English than in Chinese, and the subset of voxels where prediction accuracy is similar between the two languages (Figure S14). Results were consistent between all three subsets of voxels, suggesting that our results are not confounded by differences in prediction accuracy.

Based on Figure 4C alone, there are two possible explanations for the semantic tuning shifts. One explanation is that semantic tuning shifts are due to the specific language of the stimulus. A second explanation is that semantic tuning shifts are because of differences between native (L1) and non-native (L2) language comprehension.

To distinguish between these two possibilities, we performed a new analysis in which we compared semantic representations between two groups of participants: the L2 English speakers in our original experiment, and a separate group of L1 English speakers. Crucially, for this analysis we used brain responses to English narratives to estimate semantic representations for each group of participants. Both groups of participants read the same stimuli, and none of the stimuli were translated from a different language. For each participant we used brain responses to the English narratives to compute voxelwise semantic tuning, and projected voxelwise semantic tuning to the template space. Then, we computed group-averaged model weights for each vertex of the template space, separately for L1 speakers (β_*L*1_) and for L2 speakers (β _*L*2_ ). Finally, for each vertex we computed semantic tuning shifts between these two sets of group-averaged model weights 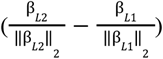, and computed the primary tuning shift index (PTSI) accordingly. Figure 4D shows vertexwise PTSI between the L1 and L2 speakers who read the same English stimuli. The PTSI shown in Figure 4D are significantly correlated with those shown in Figure 4C (r=.63; p<.001 by a one-sided permutation test). The consistency between the two sets of PTSI suggests that the semantic tuning shifts shown in Figure 4 are due to differences between L1 and L2 language comprehension, and not due to differences between the English versus Chinese stimuli, such as differences in word length, word frequency, or token rate.

There are multiple ways in which the semantic tuning shifts shown in Figure 4 could modulate semantic representations between different languages. One possibility is that these semantic tuning shifts systematically modulate semantic representations, such that voxels that are tuned towards similar word meanings shift in a consistent direction. Alternatively, there could be no correspondence between the semantic tuning of a voxel and the direction in which the semantic tuning shifts between languages. To distinguish between these two possibilities, for each voxel we used the English model weights to assign the voxel into one of the five semantic clusters shown in Figure 3A. (Results are consistent when semantic clusters are identified using Chinese model weights; Figure S16). Then we examined the direction of semantic tuning shifts for voxels in each cluster. Figure 5A shows the PTSI for each cluster and participant. For voxels in Clusters 1, 2, and 3 PTSI is positive in all participants (p<.05 for each cluster by a two-sided t-test after Fisher z-transformation). Thus, representations of family, communication, and cognition-related clusters emphasize action- and relationship-related aspects of those clusters in English and number- and collection-related aspects in Chinese. For voxels in Clusters 4 and 5 PTSI is negative in all participants (p<.05 for each cluster by a two-sided t-test after Fisher z-transformation). Thus, representations of location- and number- and name-related clusters emphasize number- and collection-related aspects in English, or action- and relationship-related aspects in Chinese.

**Figure 5.**
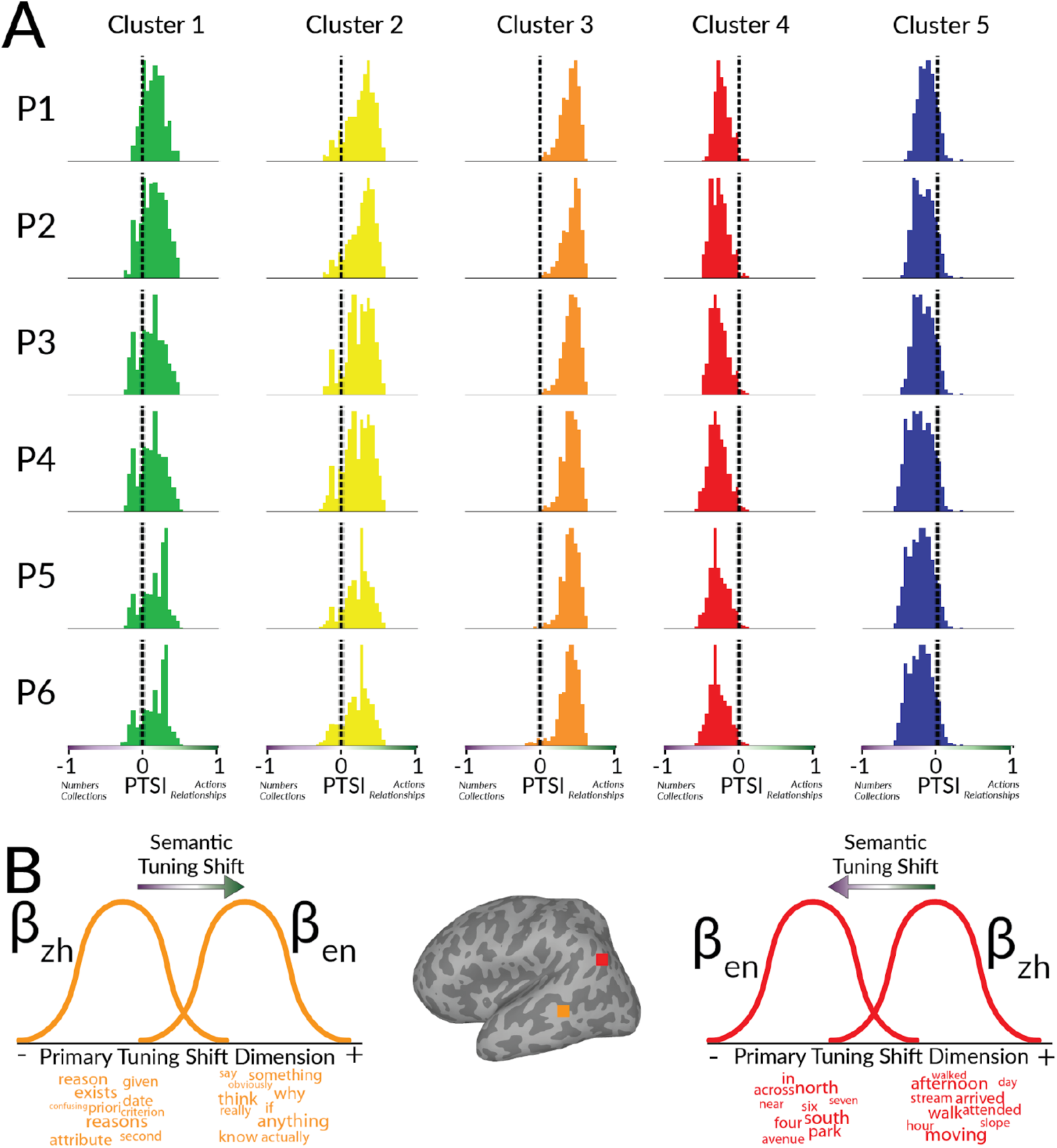
Semantic tuning shifts for different semantic clusters. Voxels were categorized into the semantic clusters shown in Figure 3. The distribution of semantic tuning shifts was examined for each semantic cluster. **A**. Semantic tuning shifts for each cluster. Histograms show the distribution of voxelwise PTSI for each cluster and participant separately. Voxels in the family-, communication-, and cognition-related clusters (Clusters 1, 2, and 3) have positive PTSI (p<.05 by a two-sided t-test after Fisher z-transformation). Thus, representations of family-, communication-, and cognition-related clusters shift to emphasize action- and relationship-related semantics in English, or number- and collection-related semantics in Chinese. Voxels in the location- and number/name-related clusters (Cluster 4 and 5) have negative PTSI (p<.05 by a two-sided t-test after Fisher z-transformation). Thus, representations of location- and name-number-related clusters shift to emphasize number- and collection-related semantics in English, or action- and relationship-related semantics in Chinese. **B**. Semantic tuning shift for two selected voxels. The voxel in frontal cortex (colored in orange) has positive PTSI. This voxel is tuned towards words related to cognition (Cluster 3) in both languages, but emphasizes action- and relationship-related aspects in English (e.g., “know”) compared to Chinese (e.g., “second”). The voxel in parietal cortex (colored in red) voxel has negative PTSI. This voxel is tuned towards words related to locations (Cluster 4) in both languages, but emphasizes number- and collection-related aspects in English (e.g., “four”) and action- and relationship-related semantics in Chinese (e.g., “moving”). These two examples illustrate how semantic tuning is modulated between languages.

To provide two specific examples of how semantic tuning shifts systematically modulate semantic representations, we show the effect of semantic tuning shifts for two selected voxels (Figure 5B). One voxel was selected from Cluster 3 and has positive PTSI. This voxel is tuned towards cognition-related word meanings in both languages, but emphasizes action- and relationship-related aspects in English (e.g., “know”) compared to Chinese (e.g., “second”). The other voxel was selected from Cluster 4 and has negative PTSI. This voxel is tuned towards location-related word meanings in both languages, but emphasizes numeric aspects in English (e.g., “four”) compared to Chinese (e.g., “moving”). These two examples show that a voxel can represent the same semantic category between languages, yet also exhibit semantic tuning shifts between languages. Overall, the results in Figure 5 show that each language systematically modulates semantic representations.

## Discussion

This study provides three unique lines of evidence for how semantic representations in the brains of bilinguals support both shared and language-specific processing between native and non-native languages. First, the same brain regions represent semantic information for both languages. Second, within these brain regions the specific voxels that are activated by a word meaning are similar between languages. Third, there are systematic, fine-grained shifts between languages in the semantic tuning of each voxel. These results are observed in each individual participant in our study, and the results generalize across participants (including the two held-out participants). Taken together, these results suggest that in the brains of bilinguals, semantic representations remain largely similar between languages, but semantic tuning is systematically modulated by each language.

Prior neuroimaging studies showed that similar brain regions are activated by different languages (14–17, 39). Understanding how semantic information is represented between different languages requires not only a comparison of brain activation, but also of the functional semantic representations within these regions. A few prior studies have compared functional representations between languages. For example, one such study compared timecourses of brain representations between participants who listened to the same narrative in different languages (20). This work reported similar brain responses between languages in regions such as the STS and middle frontal gyrus (MFG), and it reported statistically significant differences in other regions such as superior temporal gyrus (STG) and right inferior frontal gyrus (IFG). However, because that study did not explicitly model semantic representations, the differences that they observed in STG and IFG may have reflected non-semantic representations from other processes such as pragmatic inference or working memory. In contrast, our study did not observe large differences in STG and IFG. Instead, we found that semantic representations are largely similar between languages throughout the semantic system, including in STG and IFG. A different prior study explicitly modeled semantic representations in the brain (40), and showed that brain responses to sentences in one language can predict brain responses to sentences in a different language.

However, that study focused on brain responses evoked by isolated sentences, and so the results may not generalize to language comprehension in a natural context (25). Furthermore, that study did not examine differences between languages, and therefore did not show how semantic representations in the brain can encode language-specific information. Our results add to these prior studies by showing similarities and fine-grained differences in semantic representations between languages.

The bilingual participants in this study were native Chinese speakers and non-native English speakers. For this population, the two language comprehension conditions (English vs. Chinese) differed in native vs. non-native language, morphology, orthography, and language family. Thus, our findings provide strong evidence that semantic representations are similar across different languages. We find that semantic representations of the two languages are similar in many different brain regions that represent semantic information in language, including in the default mode network (41). These regions span many different timescales of language processing (42–44), which suggests that semantic representations are largely shared not only at the level of individual words, but also for more abstracted semantic representations that are built over the course of a narrative.

In this study, the encoding model prediction accuracy was much higher in Chinese (L1) than in English (L2) (Figure 2). There are many possible explanations for these differences in prediction accuracy. For example, there may have been differences between the two conditions in working memory load, high-level control, or syntactic processing, or differences between the orthography or morphology of the two languages might have affected brain activity measurements. Although all participants were fluent in both English and Chinese (Table S1), behavioral results based on self-reported comprehension responses suggest that participants found the Chinese narratives slightly easier to comprehend than they did the English ones (Figure S15). Thus, the differences in prediction accuracy might merely reflect differences in effort. Because all of the bilingual participants in our study were all L1-Chinese and L2-English, differences in prediction accuracy might be due to differences between native and non-native language processing, or differences between the stimulus languages. In future work we plan to study different bilingual populations to distinguish between these two possibilities, and to determine whether a similar difference in prediction accuracy is found in other languages and populations.

When we were analyzing the results of this study one concern was that the observed semantic tuning shifts might be an artifact of these differences in encoding model prediction accuracy. To determine if this confound could have produced our results we examined whether tuning shifts were also observed in the subset of voxels that had similar prediction accuracy across the two languages, and in the subset where prediction accuracies were higher in English than in Chinese (Figure S14). In both cases the semantic tuning shifts were still found. Thus, it appears that semantic tuning shifts are not a confound caused by differences in prediction accuracy across the two languages.

Another source of potential confounds is from the distributional word embeddings used to operationalize the meaning of each word. These embeddings reflect not only the meaning of each word, but they also reflect other word properties such as word frequency and grammatical class. We were concerned that these non-semantic word properties might have confounded our results. Therefore, we created nuisance feature spaces for word frequency and grammatical class, and re-analyzed the data with these nuisance feature spaces included. None of the conclusions of the study changed when including these additional nuisance feature spaces (Figure S17).

Thus, we do not believe that non-semantic language-related features confounded our results. However, it is impossible to account for all possible non-semantic features that may be reflected in word embeddings.

Furthermore, although there are inevitable imperfections in embedding alignment across languages (38), our tuning shift analysis assumes that embeddings are perfectly aligned across languages. Therefore it is theoretically possible that these embedding misalignments might have confounded our tuning shift results. To ensure that this was not the case we reproduced the tuning shifts across two different embedding spaces (Figures S1-S4), and also showed that the main directions of tuning shifts are different from the directions of embedding misalignment (Figure S14). Thus, the tuning shifts do not appear to be driven by embedding misalignment.

Many psycholinguistic theories explicitly differentiate between lexical and conceptual semantic representations. For example, some theories suggest that in proficient bilinguals, L1 and L2 have distinct lexical representations, but that both languages are connected to the same conceptual representations (46). Unfortunately, it is difficult to distinguish between lexical and semantic representations in our study (or indeed in any naturalistic experiment), because both of these sorts of representations are likely to be activated in the brain during natural language comprehension. Because these representations are difficult to distinguish, the semantic feature space used here likely models brain representations associated with lexical entries, and also conceptual representations retrieved from long-term memory.

The results presented here can help clarify how behavioral observations might connect to representations in the brain.Some studies observed that words in one language can semantically prime words in another language (2), and can activate translation equivalents in other languages (45). Our results show that words in one language elicit semantic representations that are similar to those in the other language, suggesting a process that could lead to semantic priming and activation of translation equivalents. Other studies observed that knowledge of one language can interfere with semantic processing in another (2–4). Our results show that semantic tuning is largely shared between languages, suggesting how learning a new language may influence semantic representations activated by the other language. Finally, behavioral studies have observed that semantic processing can differ between native and non-native languages, such as in perceived emotional intensity (6, 7). Our results show that there are systematic shifts in voxelwise semantic tuning between languages. These tuning shifts suggest how shared semantic representations can nevertheless encode word meanings distinctly for different languages.

We found that the direction of semantic tuning shifts between L1 and L2 is systematic: voxels that are tuned towards similar word meanings shift in a consistent direction, and these shifts are consistent across individuals. However, because our study only examined Chinese-English bilinguals, we cannot determine whether this effect will generalize to all L1 vs. L2 languages, or rather whether it is specific to these specific languages or to the specific cultural background of the participants. That said, two lines of evidence suggest that the direction of semantic tuning shifts is determined primarily by L1 vs. L2 comprehension. First, the most notable tuning shift changes the representation of number and action or relationship semantic concepts. This finding is consistent with prior behavioral evidence showing that semantic processing of numbers and of emotions differs between native and non-native languages, and for several different language pairs (6, 7, 9, 10, 47). Second, semantic tuning shifts between L1 and L2 comprehension were also observed within the same language (Figure 4). This consistency suggests that the tuning shifts are due to differences between L1 vs L2, rather than to the specific properties of the stimulus language. In future work, we plan to use different languages and to include broader bilingual populations in order to test how well semantic tuning shifts generalize across different languages.

Our results suggest that there are differences in how the brain represents word meaning between L1 and L2, but they do not reveal what causes these differences. We speculate that semantic tuning shifts may result from differences in the depth of language processing, working memory load, or attentional demands between native and non-native language processing. Indeed, prior studies of the human visual system have shown that differences in attentional demands may alter semantic tuning in the brain (27). Thus, differences in cognitive demands between L1 and L2 may analogously drive shifts in semantic tuning.

Our results provide the first evidence for systematic, fine-grained shifts in semantic tuning between native and non-native language comprehension. These results allow us to ground prior behavioral observations of shared and language-specific language processing in naturalistic, neurobiological evidence. The results and analysis paradigm presented here enable researchers to understand how semantic representations in the brains of bilinguals support both shared and language-specific processing.

## Methods

### Stimuli

#### Narrative transcription, translation, and preprocessing

The stimuli consisted of eleven 10- to 15 min narratives from *The Moth Radio Hour* which have been used in previous studies (18, 19, 44, 48). In each narrative, a speaker tells an autobiographical story in front of a live audience. The selected narratives cover a wide range of topics. They include both concrete and abstract words, and include words from a range of grammatical categories (See Figure S18 for details). The audio recording of each narrative was manually transcribed, and the written transcription was aligned to the audio recording. (Details of audio transcription and alignment are described in prior work (19)).

The original narratives were performed verbally in English. To construct matched Chinese stimuli, each of the English narratives was translated into Chinese by a professional translation service. To obtain word presentation times that correspond to natural speech, each translated narrative was read aloud by a professional voice actor. Then the written translations were aligned to these audio recordings. Chinese stimuli were presented with simplified Chinese characters (简化字).

#### Stimulus train and test split

Ten train narratives were used for model estimation, and one held-out test narrative was used for model evaluation. The same test narrative was used for both languages. To obtain noise-ceiling estimates, the test narrative was played to each participant four times in each language. (For P1, the test narrative was played only twice in Chinese due to a change in stimulus design after the first collected sessions.)

#### Stimulus presentation format

The words of each narrative were presented one-by-one at the center of the screen using a Rapid Serial Visual Presentation (RSVP) procedure (49). Each word was presented for a duration equal to the duration of that word in the spoken version of the narrative.

Each word was presented at the center of the screen in isolation, and a white fixation cross was present at the center of the display throughout the experiment. Participants were asked to fixate on a center cross while reading the narrative. Participants’ eye movements were monitored at 60 Hz throughout the scanning sessions using a custom-built camera system equipped with an infrared source (Avotec) and the View-Point EyeTracker software suite (Arrington Research). The eye tracker was calibrated at the end of each run of data acquisition.

Functional MRI data were collected during four 3-hour scanning sessions that were performed on different days. Each scanning session consisted of seven functional runs. Two of these runs consisted of an exact repetition of the test narrative. These repetitions were used to compute noise-ceiling estimates. The remaining five runs presented five different training narratives. The language of the training narratives was interleaved across runs.

All participants read all the narratives in both English and Chinese. Narrative presentation order was balanced between languages and across participants.

To verify that participants comprehended the narratives and paid attention to the narratives, participants were asked to recount the contents of each narrative to the experimenter at the end of each session and eye movements were manually monitored throughout the scan.

### fMRI data acquisition

Whole-brain MRI data were collected on a 3T siemens TIM trio scanner at the UC Berkeley Brain Imaging Center. A 32-channel Siemens volume coil was used. Functional scans were collected using a T2*-weighted gradient-echo EPI with repetition time (TR) 2.0045s, echo time (TE) 35ms, flip angle 74°, voxel size 2.24x2.24x4.1 mm (slice thickness 3.5mm with 18% slice gap), matrix size 100x100, and field of view 224x224 mm. Thirty axial slices were prescribed to cover the entire cortex and were scanned in interleaved order. A custom-modified bipolar water excitation radiofrequency (RF) pulse was used to prevent contamination from fat signals. Anatomical data were collected using a T1-weighted multi-echo MP-RAGE sequence on the same 3T scanner.

To stabilize head motion during scanning sessions, participants wore a personalized head case that precisely fit the shape of each participant’s head (50, 51).

### fMRI data pre-processing

Each functional run was motion-corrected using the FMRIB Linear Image Registration Tool (FLIRT) from FSL(52). All volumes in the run were averaged across time to obtain a high quality template volume. FLIRT was used to automatically align the template volume for each run to the overall template, which was chosen to be the temporal average of the first functional run for each participant. These automatic alignments were manually checked and adjusted as necessary to improve accuracy. The cross-run transformation matrix was then concatenated to the motion-correction transformation matrices obtained using MCFLIRT, and the concatenated transformation was used to resample the original data directly into the overall template space. Noise from motion, respiratory, and cardiac signals were removed with a component-based detrending method (CompCor (53)). Responses were z-scored separately for each voxel and narrative. During z-scoring, the mean response across time was subtracted and the remaining response was scaled to have unit variance. Before data analysis, 10 TRs from the beginning and 10 TRs at the end of each narrative were discarded in order to account for the 10 seconds of silence at the beginning and end of each scan and to account for non-stationarity in brain responses at the beginning and end of each scan.

### Cortical surface reconstruction and visualization

Cortical surface meshes were generated from the T1-weighted anatomical scans using FreeSurfer software^32^. Before surface reconstruction, anatomical surface segmentations were carefully hand-checked and corrected using Blender software and pycortex (54, 55). Relaxation cuts were made into the surface of each hemisphere. Blender and pycortex were used to remove the surface crossing the corpus callosum. The calcarine sulcus cut was made at the horizontal meridian in V1 using retinotopic mapping data as a guide.

Functional images were aligned to the cortical surface using pycortex. Functional data were projected onto the surface for visualization and analysis using the line-nearest scheme in pycortex. This projection scheme samples the functional data at 32 evenly spaced intervals between the inner (white matter) and outer (pial) surfaces of the cortex and then averages together the samples. Samples are taken using nearest-neighbor interpolation, wherein each sample is given the value of its enclosing voxels.

### Cortical parcellation

FreeSurfer ROIs were used to anatomically localize the temporal, parietal, and prefrontal regions for each participant. ROIs were based on the Desikan-Killiany atlas (32). The ROIs used for the temporal region were *“bankssts”, “inferiortemporal”, “middletemporal”, “superiortemporal”, “temporalpole”, “transversetemporal”, “fusiform”, “entorhinal”, “parahippocampal”*. The ROIs used for the parietal region were *“inferiorparietal”, “superiorparietal”, “supramarginal”, “precuneus”, “isthmuscingulate”, “posteriorcingulate”*. The ROIs used for the prefrontal region were *“caudalmiddlefrontal”, “parsopercularis”, “parsorbitalis”, “parstriangularis”, “rostralmiddlefrontal”, “superiorfrontal”, “frontalpole”, “caudalanteriorcingulate”*.

### Localizers for known ROIs

Known ROIs were localized separately in each participant using a visual category localizer and a retinotopic localizer (56, 57). Details of localizer experiments are provided in prior work (18, 19).

### Participants

Functional data were collected from six bilingual participants who were fluent in both Mandarin Chinese (native) and English (non-native): P1 (29F), P2 (25M), P3 (25F), P4 (25M), P5 (24F), P6 (26M). Participant P1 was an author of this paper. Language proficiency of each participant was evaluated by the Language Experience and Proficiency Questionnaire (LEAP-Q) (58) and the Language History Questionnaire (LHQ3) (59). Participants began English language acquisition between the ages of 2 and 11, and spent between 5 and 12 years in a country where English is spoken. At the time of the experiment, participants primarily used Chinese in interactions with family, English in interactions at school/work, and a mix of the two languages in interactions with friends. Please see Table S1 for additional details of participants’ use of each language.

All participants were healthy and had normal or corrected-to-normal vision. All subjects were right handed or ambidextrous according to the Edinburgh handedness inventory (laterality quotient of -100: entirely left-handed, +100: entirely right-handed) (60). Laterality scores were +5, 0, +90, +100, +65, +65 for P1-6 respectively.

A separate set of functional data were collected from six participants between the ages of 24 and 31 who were native English speakers. Data for P7-P12 were originally collected for separate studies (19, 61).

### Statistical analysis

Voxelwise modeling (VM) was used to model BOLD responses (62, 63). In the VM framework, stimulus and task parameters are nonlinearly transformed into sets of features (also called feature spaces) that are hypothesized to be represented in brain responses. Linearized regression is used to estimate a separate encoding model for each voxel and feature space. Each encoding model describes how a feature space is represented in the BOLD response of a voxel. A held-out dataset that was not used for model estimation is used to evaluate model prediction accuracy and to determine the significance of the model prediction accuracy. Encoding models were estimated and evaluated separately for each participant.

All model fitting and analysis was performed using custom software written in Python, making heavy use of NumPy (64), SciPy (65), Matplotlib (66), Himalaya (67), and Pycortex (55).

#### Construction of semantic feature spaces

To capture the semantic content of the stimulus narratives, each word of the stimulus narrative was projected to a 300-dimensional embedding space (28). Embedding spaces were constructed separately for English and for Chinese. The embedding spaces were orthogonally transformed to align the English and Chinese embedding spaces, such that there is a high correlation between embeddings of words in different languages that have similar meanings (29). This orthogonal transformation was necessary to ensure that reported similarities and differences in model weights are indeed due to underlying brain responses, rather than differences between the embedding space of each language. Furthermore, the estimated semantic models are invariant to orthogonal rotation of the feature spaces because encoding models linearly map from the feature space to brain responses. Given a (num_embedding_dimensions x num_timesteps) feature space matrix *X* and a (num_timesteps) voxel response vector *y*, we estimate a (num_embedding_dimensions) vector of weights β such that *y*≈β*X*. The estimated weights β reflect the semantic tuning of the voxel: the estimated activation of the voxel in response to a word with embedding *v* is β^*T*^ *v*. If *X* is orthogonally rotated with the matrix *M*, the estimated weights would become *M*β. But the embedding of each word would also have been rotated by *M*, so the orthogonal rotation of the feature space would not change the estimated tuning of each voxel: with the rotated embedding space, the estimated activation in response to the same word would be (*M*β)^*T*^ (*Mv*) = β^*T*^ *M*^*T*^ *Mv* = β^*T*^ *v*, which was the same as the model estimated before orthogonally rotating the embedding space.

Because of non-isometry between embedding spaces of different languages and limitations to multilingual alignment procedures (38), embedding spaces are not perfectly aligned between languages (e.g., “table” and “桌子” [table] may not project to exactly the same vector). To ensure that results are not specific to the idiosyncrasies of a particular embedding space or a particular imperfection in cross-lingual embedding alignments, all analyses were replicated with a different embedding space, multilingual BERT (mBERT, *bert-base-multilingual-cased (37)*). mBERT is a twelve-layer contextual language model that was jointly trained on text from 104 languages. No explicit cross-lingual alignment object was included during the training of mBERT, but embeddings in some layers are implicitly aligned over the course of training (68). To obtain word embeddings from mBERT, each sentence of the stimulus narratives was provided as input to mBERT and then the 768-dimensional activation of layer nine was used as an embedding of each word. Layer 9 was chosen because it produces the best aligned embeddings (Figure S19), and because intermediate layers of contextual language models have been shown to produce the most accurate predictions of brain responses (44, 69–72).

#### Construction of low-level feature spaces

Seven low-level feature spaces were constructed to account for the effects of low-level stimulus information on BOLD responses. For both languages, models were fit with feature spaces that reflect visual spatial and motion features (motion energy) (73–75), word count, single phonemes, diphones, triphones (76), and intermediate level features that capture orthographic similarities by measuring the pixelwise overlap between words (77). For English, an additional low-level feature space reflected letter count. For Chinese, an additional low-level feature space reflected character count.

#### Stimulus feature space preprocessing

Before voxelwise modeling, each stimulus feature was truncated, downsampled, z-scored, and delayed. Data for the first 10 TRs and the last 10 TRs of each scan were truncated to account for the 10 seconds of silence at the beginning and end of each scan and to account for non-stationarity in brain responses at the beginning and end of each scan. An anti-aliasing, 3-lobe Lanczos filter with cut-off frequency set to the fMRI Nyquist rate (0.25 Hz) was used to resample the stimulus features to match the sampling rate of the fMRI recordings. Then the stimulus features were each z-scored in order to account for z-scoring performed on the MRI data (For details see *MRI data collection*). In the z-scoring procedure, the value of each feature channel was separately normalized by subtracting the mean value of the feature channel across time and then dividing by the standard deviation of the feature channel across time. Lastly, finite impulse response (FIR) temporal filters were used to delay the features in order to model the hemodynamic response function of each voxel. The FIR filters were implemented by concatenating feature vectors that had been delayed by 2, 4, 6, and 8 seconds (following prior work(18, 19, 78)). A separate FIR filter was fit for each feature, participant, and language.

#### Voxelwise encoding model fitting

Voxelwise encoding models were estimated in order to determine which features are represented in each voxel. Each model consists of a set of regression weights that describes BOLD responses in a single voxel as a linear combination of the features in a particular feature space. Regression weights were estimated using banded ridge regression(79). Unlike standard ridge regression, which assigns the same regularization parameter to all feature spaces, banded ridge regression assigns a separate regularization hyperparameter to each feature space. Banded ridge regression thereby avoids biases in estimated model weights that could otherwise be caused by differences in feature space distributions. Mathematically, for a train dataset with *v* voxels and *n* TRs, the *m* delayed feature spaces *F* _*i*_ (*X*), *i*∈{1, …, *m*} (each dimension *p*_*i*_ ) were concatenated to form a feature matrix *F*’(*X*) (dimension 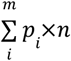). Then banded ridge regression was used to estimate a mapping *B* (dimension 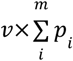) from *F*’(*X*) to the matrix of voxel responses *Y* (dimension *v*×*n*). *B* is estimated according to 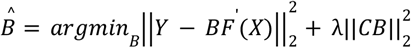. The diagonal matrix *C* of regularization hyperparameters for each feature space and each voxel is optimized over 10-fold cross-validation. See *Regularization hyperparameter selection* for details.

#### Stepwise regression procedure

To remove confounds from stimulus correlations between semantics and low-level sensory stimulus features, a stepwise regression procedure was used. First banded ridge regression was used to jointly estimate encoding models that predict BOLD responses from the seven low-level stimulus features. Only data from the train narratives was used to estimate models. Then, the low-level models were used to predict BOLD responses to the train and test narratives. The predicted BOLD responses *Ŷ* _*lowlevel,train*_ and *Ŷ* _*lowlevel,test*_ were subtracted from the true BOLD responses *Y* _*train*_, *Y* _*test*_ . The residual BOLD responses *Y* _*train*_ − *Ŷ* _*lowlevel,train*_, *Y* _*test*_ − *Ŷ* _*lowlevel,test*_were zscored. During z-scoring, for each voxel separately the mean response across time was subtracted and the remaining response was scaled to have unit variance. The z-scored residual BOLD responses were used to estimate encoding models that predict BOLD responses from semantic stimulus features.

#### Regularization hyperparameter selection

Five-fold cross-validation was used to find the optimal regularization hyperparameters for each feature space and voxel. Hyperparameter candidates were chosen with a random search procedure (80): 1000 normalized hyperparameter candidates were randomly sampled from a dirichlet distribution and were then scaled by 21 log-spaced values ranging from 10^−10^ to 10^10^. The regularization hyperparameters for each feature space and voxel were selected as the hyperparameters that produced the minimum squared error (L2) loss between the predicted voxel responses and the recorded voxel responses 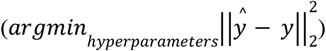. Regularization hyperparameters were chosen separately for each participant and language. Hyperparameter search was performed using the Himalaya Python package (67).

#### Model estimation and evaluation

The selected regularization hyperparameters were used to estimate model weights that map from the semantic feature space to voxel BOLD responses. Model weights were estimated separately for each voxel, language, and participant. The model weights for each voxel and language reflect the semantic tuning of the voxel in that language.

The test dataset was not used to select hyperparameters or to estimate regression weights. The prediction accuracy *R*^2^of the feature spaces was computed per voxel as the coefficient of determination between the predicted voxel responses and the recorded voxel responses on the test dataset. To determine which voxels represent semantic information in each language, prediction accuracy was computed for within-language predictions (train and test on the same language). To determine how well model weights estimated for one language generalize to the other language, prediction accuracy was also computed for across-language predictions (train on one language, and test on a different language).

A permutation test with 1000 iterations was used to compute the statistical significance of prediction accuracy. In each permutation, the test responses were shuffled in blocks of 10 TRs (19, 44, 71, 81–85). Shuffling was performed in blocks of 10 TRs in order to preserve autocorrelations in voxel responses. Then the prediction accuracy ( *R*^2^) was computed between the predicted responses and the permuted test responses. The distribution of test accuracies over permutation iterations was used as a null distribution to compute the p-value of prediction accuracy for each voxel. A Benjamini-Hochberg correction for multiple comparisons was applied to the voxelwise p-values (86). Permutation tests were performed separately for each voxel, language, and participant.

Noise-ceiling correction was performed by normalizing the prediction accuracy of each voxel *R*^2^ by the maximum possible prediction accuracy (*noise-ceiling*) (33–35). To compute the noise-ceiling, first the maximum explainable variance (*EV*, also referred to as *signal power*) is computed for each voxel. EV measures the consistency of measured BOLD responses over repeated stimulus presentations, and reflects the amount of response variance in the test data that could be explained by a perfect model. Formally, for a test dataset with *N* repeats of a *T* TR test narrative, and recorded BOLD responses *y*_1_ … *y* _*N*_ ∈ *R*^*T*^ for a single voxel, EV is defined as follows (each *y* is first zscored across time):

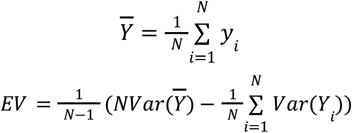

The noise-ceiling 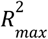 is obtained by dividing the EV of each voxel by 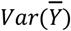. The noise-ceiling corrected prediction accuracy 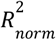 is then obtained for each voxel by dividing the prediction accuracy *R*^2^ by the noise-ceiling 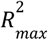 . For very noisy voxels, the estimated noise-ceiling may be lower than the measured *R*^2^ and therefore lead to divergent estimates of 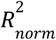. We used a heuristic to correct for this divergence. We identified the set of voxels where *R*^2^ is greater than 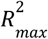, selected the maximum 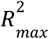 over these voxels, and then clipped 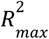 to be above this maximum value. Note that this heuristic results in a conservative estimate of the noise-ceiling corrected prediction accuracy.

#### Group-level prediction accuracy

Group-level prediction accuracy was computed by computing prediction accuracy for each participant in the participant’s native brain space, and then projecting individual participant results into a template space (fsAverage (30)). Average prediction accuracy across six participants was computed for each fsAverage vertex.

#### Generalization to new participants

To ensure generalization to new participants, two steps were performed. First, the entire analysis was performed at the individual participant level – group-averaged results are shown only as summary statistics. Second, before the final analyses were performed, two out of the six participants were set aside as held-out participants. The data for these participants were not analyzed until the data analysis and interpretation pipeline was finalized (87).

### Voxel Selection for Tuning Analyses

To ensure that semantic tuning shift analyses were performed only on voxels that represent semantic information in both English and Chinese, all of the following model weight interpretation analyses were performed only on voxels that were well-predicted ( √*R*^2^ > 0. 1) in both English and Chinese.

### Semantic Tuning Shifts

The semantic tuning shift of each voxel was used to describe how voxelwise semantic tuning changes between languages. First, the model weights were normalized for each language and voxel by dividing each 300-dimensional vector of model weights by the L2-norm of the vector. Then, the semantic tuning shift of each voxel was computed by subtracting the normalized Chinese model weights from the normalized English model weights. Formally, the semantic tuning shift for a voxel with weights β_*en*_ and β_*zh*_ was defined as 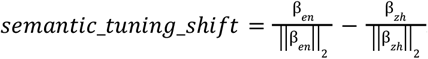. Results were qualitatively similar when L1 normalization was used instead of L2 normalization, and when normalization was performed after subtracting the model weights between the two languages (Figure S20).

The semantic tuning shift vector for each voxel describes how semantic tuning changes from Chinese to English. For example, for a voxel that becomes more tuned towards number-related semantics when the stimulus language changes from Chinese to English, the semantic tuning shift vector would point in the direction of number semantics in the embedding space.

Note that defining the semantic tuning shift as the change in tuning from Chinese to English 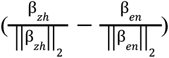 would change the sign of the semantic tuning shift vector but also swap the languages on each end of the vector. Thus, the choice to define semantic tuning shift as the shift from Chinese to English rather than from English to Chinese does not affect the reported results.

#### Dimensions of semantic tuning shift

Principal component analysis (PCA) was used to determine the main directions of semantic tuning shifts. Because model weights accurately describe semantic tuning only for well-predicted voxels, we only investigated semantic tuning shifts for voxels that were well-predicted in both languages (see *Voxel Selection for Tuning Analyses* for details; selecting voxels based on significance instead of prediction accuracy produces similar results as shown in Figure S21). To increase the influence of better-predicted voxels on the estimated principal components (PCs), the semantic tuning shift of each voxel was scaled by the voxel’s mean prediction accuracy between languages. The semantic tuning shift vectors were concatenated across participants and languages. PCA was applied to the concatenated semantic tuning shift vectors to find a set of orthogonal axes that best explain variance in voxelwise semantic tuning shift vectors. The PCs that explain the highest variance in voxelwise semantic tuning shift describe the primary semantic dimensions of semantic tuning shifts. We refer to the PC that explains the most variance in voxelwise semantic tuning shifts as the *first semantic tuning shift PC*. To examine whether the semantic tuning shift for each voxel is towards one end of the first semantic tuning shift PC or the other, the Pearson correlation was computed between the semantic tuning shift vector of each voxel and the PC. For each voxel we refer to this correlation as the *primary tuning shift index* (PTSI).

### Weight Clustering

A clustering approach was used to separate voxels into groups that have similar semantic tuning^40^. Model weights for each participant and language were projected to a standard template space (fsAverage^32^). Each projected model weight β _*vertex,participant,language*_ ∈ *R*^300^ was normalized to have unit L2-norm: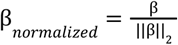. The normalized model weights were averaged across participants and languages, and then hierarchical clustering was performed on the normalized group-averaged model weights. Clustering was only performed on vertices that were well-predicted in both languages (group-averaged 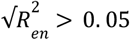 and 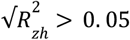). In total, there are 12 sets of model weights across the six participants and two languages. The number of clusters was chosen based on a leave-one-out cross-validation procedure. One of the twelve sets of model weights was held out in each cross-validation fold. The remaining eleven sets of model weights were averaged across participants and languages for each vertex of the template space, and hierarchical clustering was used to obtain *N* group-level clusters, separately for *N* ranging from 2 through 25. For each number *N* of clusters, the group-level model weight clusters were used to predict BOLD responses for the held out participant and language in the template space. The cross-validation accuracy was computed for each vertex as the coefficient of determination (*R*^2^) between the predicted and recorded BOLD responses. Only vertices that were well-predicted in both languages (group-averaged 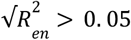 and 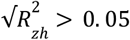) were included in this analysis. The cross-validation score plateaus around five clusters (Figure S9). Thus, we chose to use five clusters for the analyses shown in Figure 3 and Figure 5. This clustering procedure resulted in five 300-dimensional cluster centroids. These centroids define clusters of voxels that are each tuned towards similar word meanings. Voxels were assigned to clusters based on the Pearson correlation between voxelwise model weights and each of the 300-dimensional cluster centroids. Cluster assignments were performed separately for each participant and language.

### Semantic Feature Space Interpretation

The semantic cluster centroids are points within the word embedding space. To qualitatively interpret these centroids, the closest stimulus words were identified. For each centroid, the closest stimulus words were defined as the words with embeddings that have the largest Pearson correlation with the vector that corresponds to that centroid. For all word clouds, we removed emotionally charged words that could be distressing to some readers. The same procedure was used to create word clouds used to interpret the first semantic tuning shift PC.

## Acknowledgments

This work was supported by the following grant awards: National Science Foundation (NSF Nat-1912373); German Federal Ministry of Education and Research (BMBF 01GQ1906); Berliner ChancengleichheitsProgramm (BCP) to FD; a National Science Foundation GRFP (DGE 1752814) to CC; an IBM PhD Fellowship to CC; and by internal UC Berkeley funds. Portions of this work were developed from the doctoral thesis of CC. We thank Tom Dupré la Tour, Anwar Nunez-Elizalde, Matteo Visconti di Oleggio Castello, and Mathis Lamarre for their help in various stages of this project. We thank Cong Du for their help with stimulus generation.

## Author contributions

CC, XG, CT, JG and FD conceived and designed the experiments. CT and XG recruited bilingual participants. CC, CT, and XG collected fMRI data. XG generated the stimuli, preprocessed the fMRI data, and performed surface segmentation. CC performed analyses. CC and FD wrote the manuscript. All authors contributed to editing the manuscript. FD and JG directed and supervised all aspects of this project.

## Data availability

Data will be made publically available on GIN (https://gin.g-node.org) immediately upon acceptance.

## Code availability

Analysis code will be made publically available immediately upon acceptance.

