## Supplemental Information for "Bilingual language processing relies on shared semantic representations that are modulated by each language"

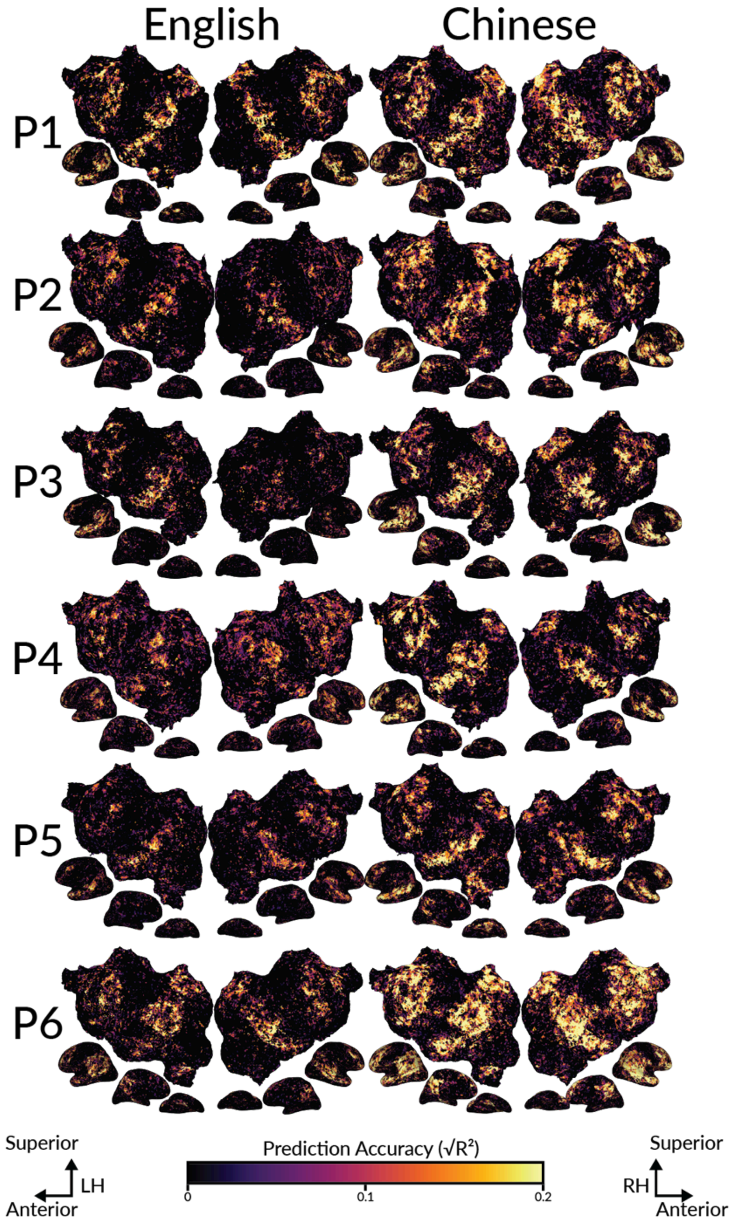


**Figure S1. Cortical distribution of semantic representations for each language when mBERT is used as the semantic feature space.** To validate the results shown in Figure 2 multilingual BERT (mBERT) was used instead of fastText as a semantic feature space. VM was used to estimate model weights that map from mBERT features to BOLD responses in each voxel and for each language separately. Estimated model weights for each language were used to predict voxelwise BOLD responses to a held-out dataset in the same language. Prediction accuracy was computed as the CD ($R^2$) between predicted and recorded BOLD responses. Prediction accuracy for each participant and language is shown on the flattened cortical surface of the participant's native brain space. For both languages and in each participant the highest prediction accuracy (brightest voxels) is found within the bilateral temporal, parietal, and prefrontal cortices. This suggests that the same brain regions are well-predicted for both languages and that these results are not dependent on the specific semantic feature space that is used.

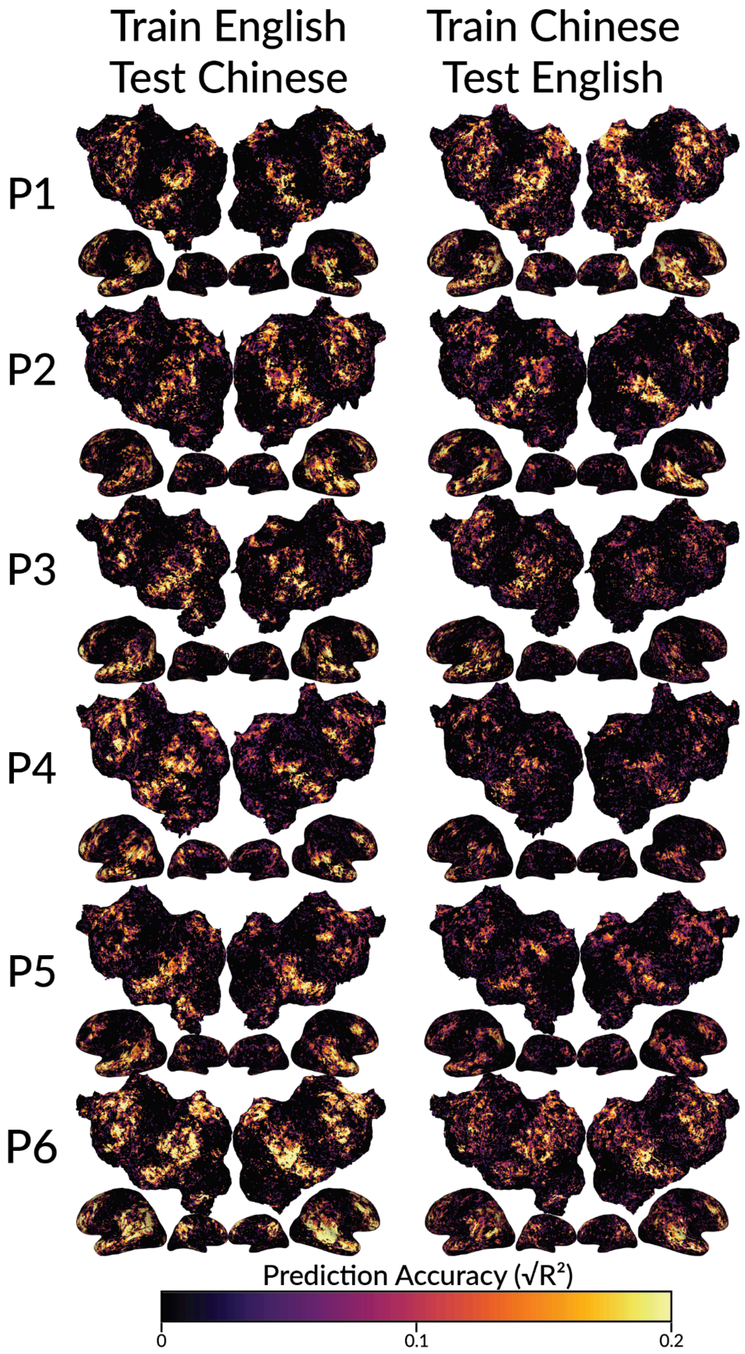


**Figure S2: Shared semantic representations between languages as shown by across-language prediction accuracy when mBERT is used as the semantic feature space.** Across-language prediction accuracy is shown for each participant on the flattened cortical surface of the participant's native brain space. Estimated voxelwise model weights in one language were used to predict the held-out dataset in the other language. Prediction accuracy was computed as the CD ($R^2$) between predicted and recorded BOLD responses. Prediction accuracy is given by the color scale. Well-predicted voxels appear brighter. In each participant, the semantic model estimated for one language accurately predicts voxel responses to the other language throughout the semantic system. Thus, semantic representations within the semantic system are largely shared between languages.

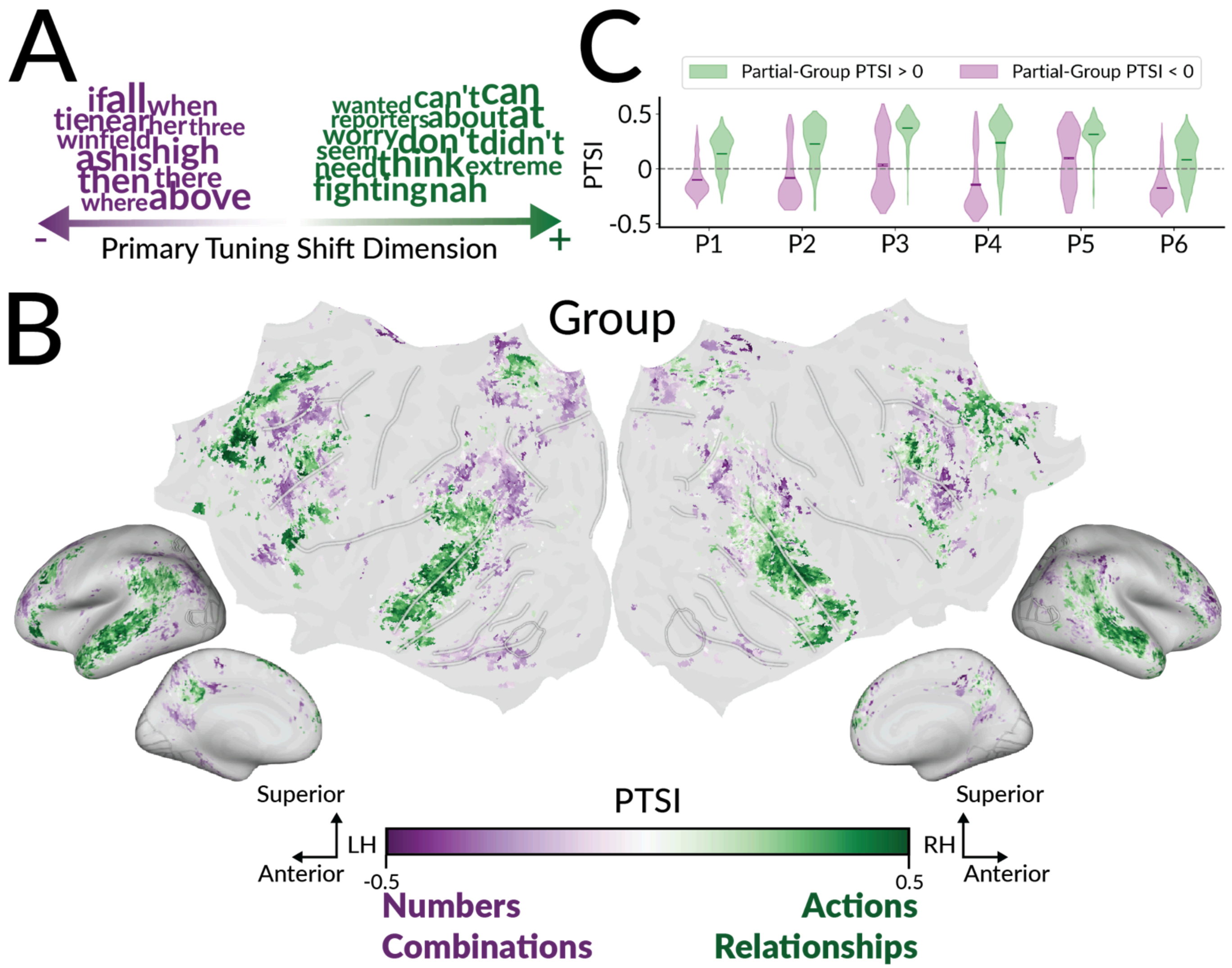


**Figure S3: Cortical distribution of semantic tuning shifts between languages when mBERT is used as the semantic feature space.** To validate the results in Figure 4 multilingual BERT (mBERT) was used as the semantic feature space. The semantic tuning shift of each voxel was defined as the change in model weights between English and Chinese ($semantic\_tuning\_shift = \frac{\beta_{en}}{\|\beta_{en}\|_2} - \frac{\beta_{zh}}{\|\beta_{zh}\|_2}$). For each voxel the semantic tuning shift describes which concepts elicit higher BOLD responses in one language relative to the other. Principal component analysis (PCA) was used to determine the main dimensions of voxelwise semantic tuning shifts.

**A.** Interpretation of the first semantic tuning shift PC. To identify words that were closest to each end of the PC, we computed the Pearson correlation between the contextual embedding of each word in the English stimulus and the PC. Naively comparing the contextual embedding of each stimulus word is heavily biased towards very frequent words, because the same word can have multiple contextual embeddings. Thus the top 1% of most frequent words were removed from this analysis. Words closest to one end of the PC are shown in purple. Words closest to the other end of the PC are shown in green. Semantic tuning shifts towards one end of the PC emphasize number/direction-related semantics (purple), while semantic tuning shifts towards the other end of the PC emphasize action-related semantics (green). The interpretation of the PC is broadly consistent with the results shown in Figure 4A for the fastText semantic feature space.

**B**. The Pearson correlation between the semantic tuning shift vector and the PC was computed. We refer to this correlation as the *primary tuning shift index* (PTSI). PTSI is shown on the flattened cortical surface of the template space. Vertices shown in purple shift towards one end of the PC. Vertices shown in green shift toward the other end of the PC. Vertices that were not well-predicted in both languages ($\sqrt{R^2} > 0.1$) in at least one

participant are shown in grey. The cortical distribution of PTSI is consistent with the results shown in Figure 4B for the fastText model.
**C**. Consistency in the cortical distribution of PTSI between each participant and the rest of the group. For each participant, the other five participants were used to compute a partial-group estimate of PTSI for each vertex. For each participant, green violin plot depicts the distribution of PTSI over vertices in which partial-group PTSI is positive, and the purple violin plot depicts the distributions PTSI over vertices in which partial-group PTSI is negative. For each participant vertices with positive PTSI tend to also have positive PTSI in the partial-group ($p<.05$ by a one-sided t-test after Fisher z-transformation, except P3 and P5) and vertices with negative PTSI tend to also have negative PTSI in the partial-group ($p<.05$ by a one-sided t-test after Fisher z-transformation). Thus, the cortical distribution of PTSI is consistent between participants. Overall, there are systematic semantic tuning shifts between languages that are consistent across participants. These results suggest that semantic representations are modulated by each language.

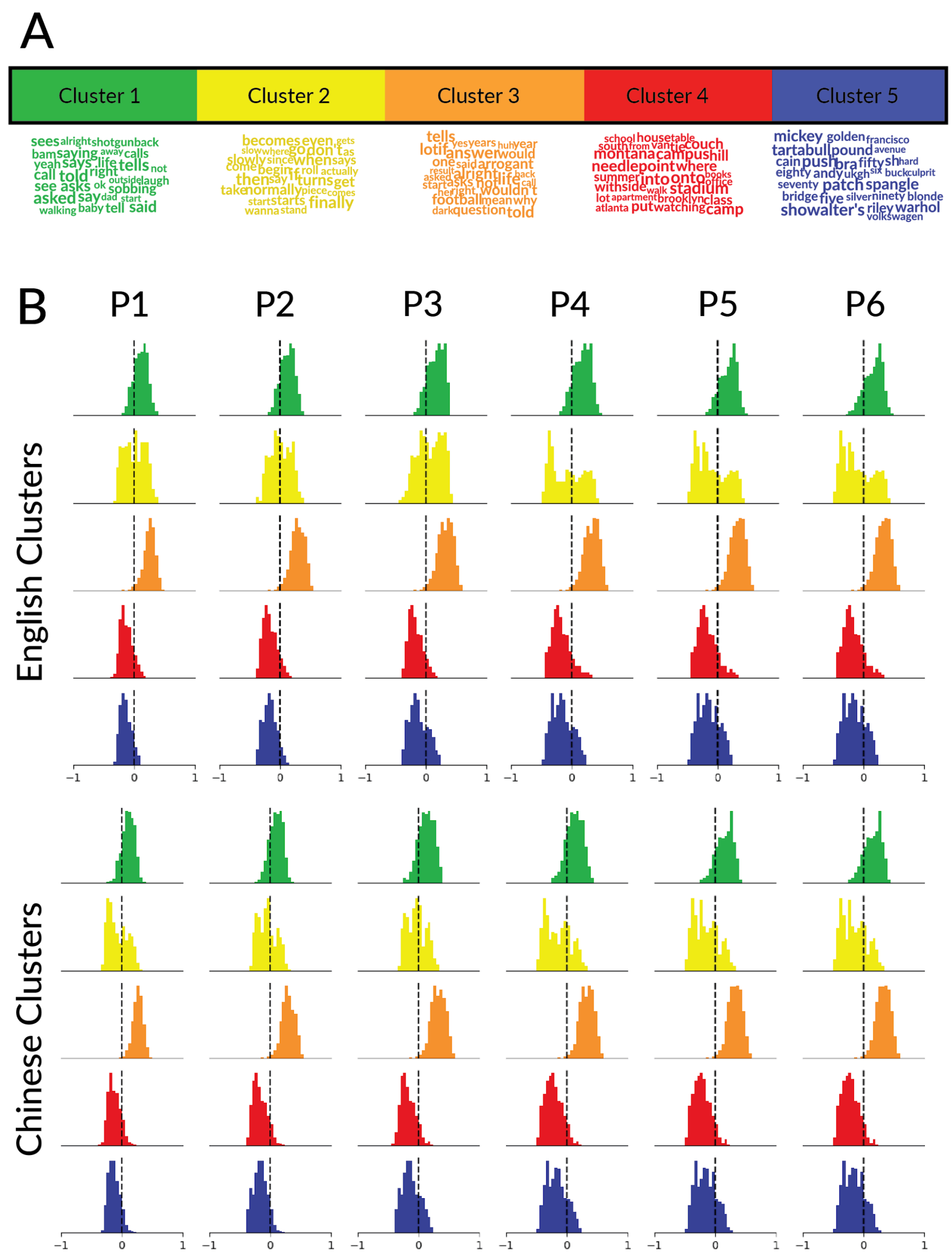


**Figure S4: Semantic tuning shifts per semantic cluster when mBERT is used as the semantic feature space.** To validate the results in Figure 5 multilingual BERT (mBERT) was used as the semantic feature space in VM. For each participant and language separately, model weights were used to assign each voxel to one of five semantic clusters. **A.** The meaning of each cluster was determined by finding the stimulus words closest to the cluster centroid. The clusters represent semantics related to communication (Cluster 1, green), cognition (Cluster 3, orange), locations (Cluster 4, red), and numbers/names (Cluster 5, blue). One cluster (Cluster 2, yellow) represents miscellaneous adjectives. Clusters are semantically noisier than for fastText (Figure 3A), likely because contextual mBERT embeddings capture more syntactic information than lexical embeddings such as fastText. **B**. Voxels were assigned to clusters based on English or Chinese mBERT model weights. The direction of semantic tuning shift of each voxel in each cluster was computed using the PTSI metric which is the Pearson correlation between the semantic tuning shift of the voxel and the first semantic tuning shift PC (as in Figure 4). Histograms indicate the distribution of PTSI values for voxels in each cluster and participant separately. Semantic tuning shifts for voxels in the communication-, and cognition-related clusters (Clusters 1 and 3) have positive PTSI. Thus, representations of communication-, and cognition-related concepts shift to emphasize action/relationship-related semantics in English as compared to Chinese. In contrast, semantic tuning shifts for voxels in the location- and number/name-related clusters (Cluster 4 and 5) have negative PTSI.

Semantic tuning shifts for voxels in Cluster 2 (miscellaneous adjectives) are mixed. Overall, representations of location- and number/name-related concepts shift to emphasize number/collection-related semantics in English as compared to Chinese. These results suggest that voxels that represent similar semantic concepts shift in similar directions between languages.

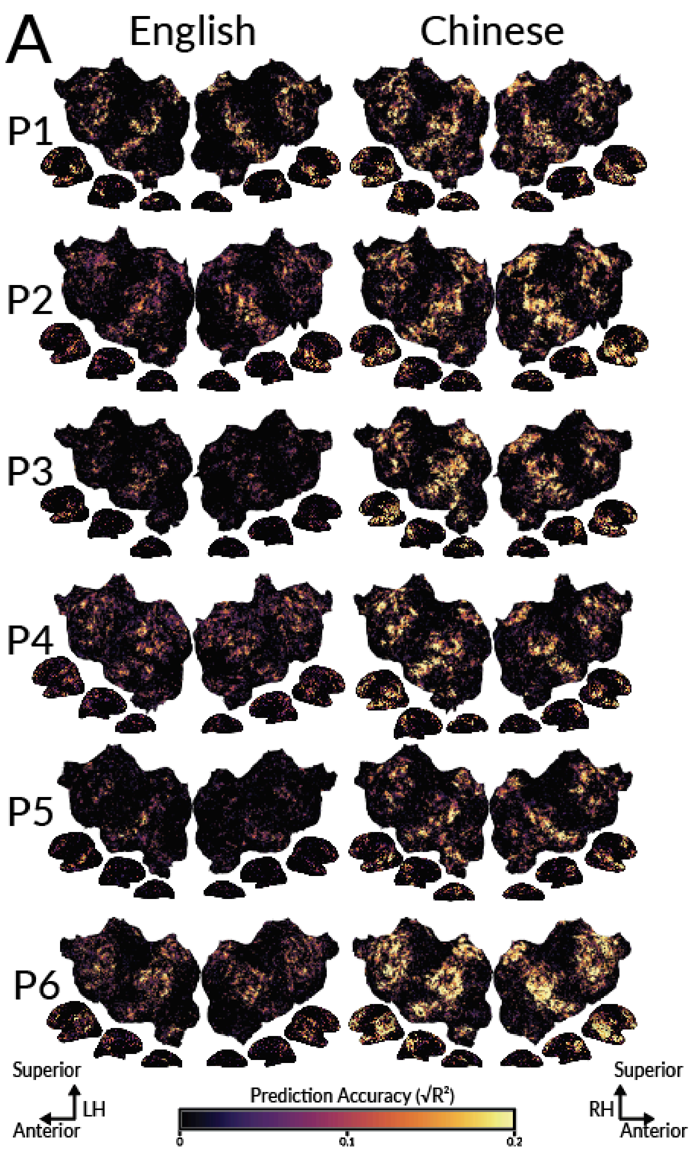


**Figure S5. Cortical distribution of semantic representations for each language and in each individual participant.** To determine where semantic information is represented for each language, voxelwise models estimated for each language were used to predict held-out data for the same language (within-language prediction accuracy). FastText was used as the semantic feature space. Prediction accuracy was computed as the CD ($R^2$) between predicted and recorded BOLD responses. Prediction accuracy is shown for each language and participant separately. Results are shown on the flattened cortical surface of each participant's native brain space. The color of each vertex indicates prediction accuracy according to the colorbar at the bottom. The magnitude of prediction accuracy differs between participants, partially reflecting individual differences in signal quality (Supplementary Figure S7). For both languages the highest prediction accuracy (brightest voxels) is found within the bilateral temporal, parietal, and prefrontal cortices. Prediction accuracy is significantly positively correlated between languages in each participant (r=0.49, 0.36, 0.28, 0.32, 0.19, 0.46 for S1-S6; one-sided p<.05 for each participant by a permutation test). This suggests that the same brain regions within the semantic system are well-predicted for both languages.

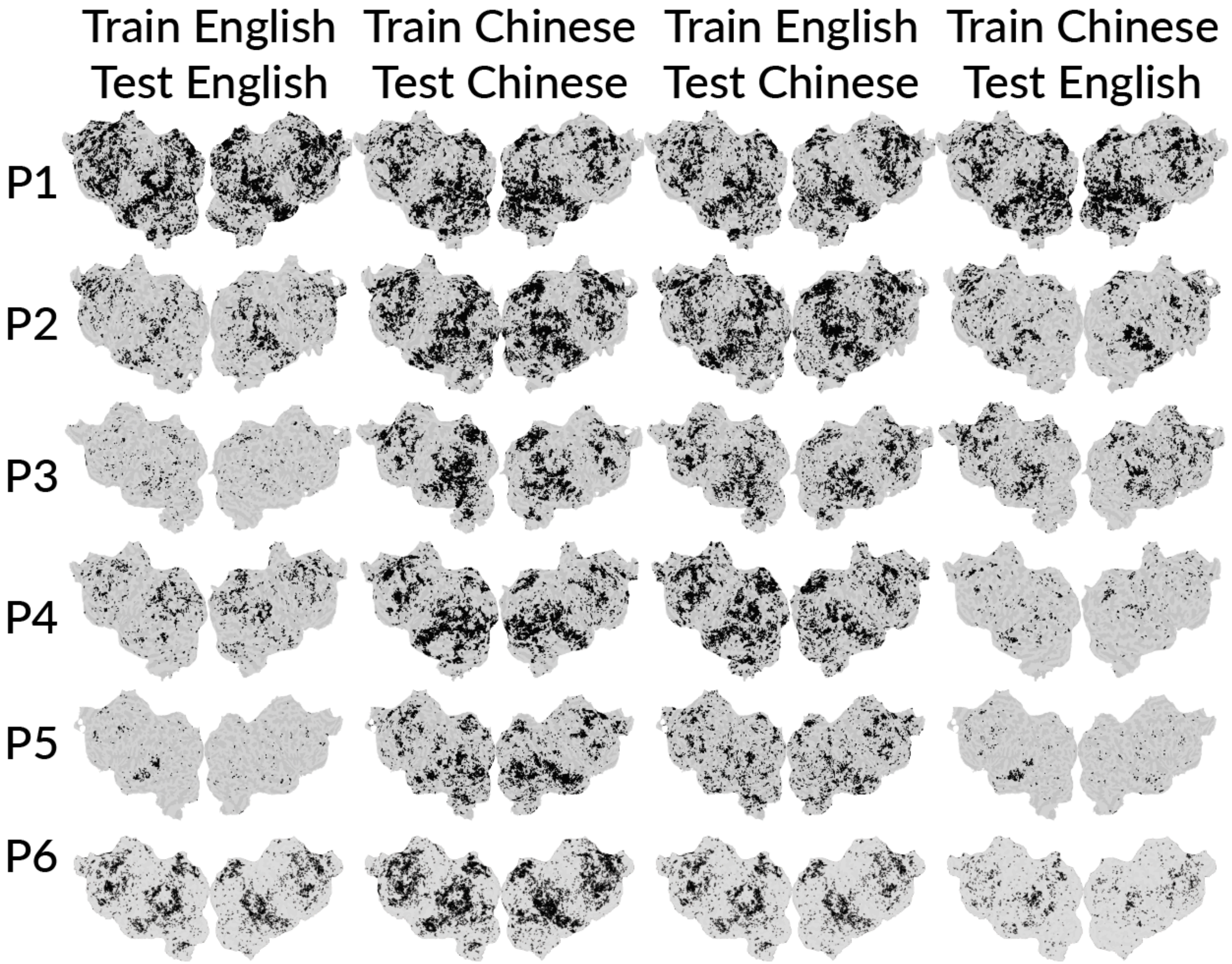


**Figure S6: Statistical significance of prediction accuracy for each participant and language.** Estimated model weights for the fastText semantic feature space were used to predict voxel responses on a held-out dataset. Within-language and across-language prediction accuracy were computed for each voxel as the CD ($R^2$) between predicted and true BOLD responses on the held-out dataset. The statistical significance of prediction accuracy for each voxel was determined by comparing the prediction accuracy of the estimated models to the prediction accuracy in predicting permuted data. The set of voxels that were significantly well-predicted are shown on the flattened cortical surface of each participant, separately for each language, and separately for within- and across-language testing. Voxels shown in black were significantly well-predicted (one-sided $p<.05$, FDR corrected with a Benjamini-Hochberg correction for multiple comparisons). The number of significantly well-predicted voxels differs across participants, reflecting individual differences in prediction accuracy shown in Supplementary Figures S5 and S11. Across participants, significantly well-predicted voxels are found within the semantic system, both within- and across-languages.

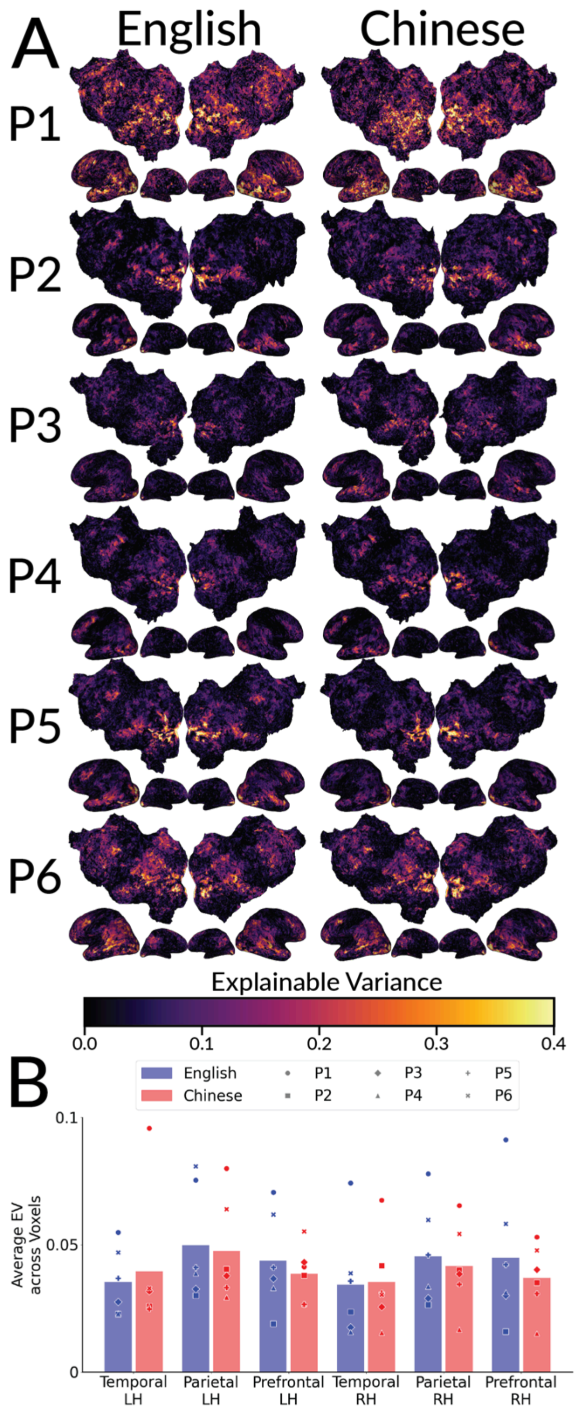


**Figure S7: Explainable variance for each participant and language.**
Explainable Variance (EV) was computed as a measure of the noise-ceiling for each participant, voxel and language separately.
**A**. The explainable variance (EV) of each voxel is shown on the flattened cortical surface of each participant's native brain space. EV is shown for English and Chinese separately. The color of each voxel indicates explainable variance according to the colorbar at the bottom. The magnitude of EV varies across participants, suggesting individual differences in the quality of the recorded BOLD signal.
**B**. For each brain region in the semantic system, the average EV across voxels is shown for each participant and language separately. Blue markers indicate EV in English, and red markers indicate EV in Chinese. Bars indicate the mean across participants. EV is not consistently higher in one language than the other.

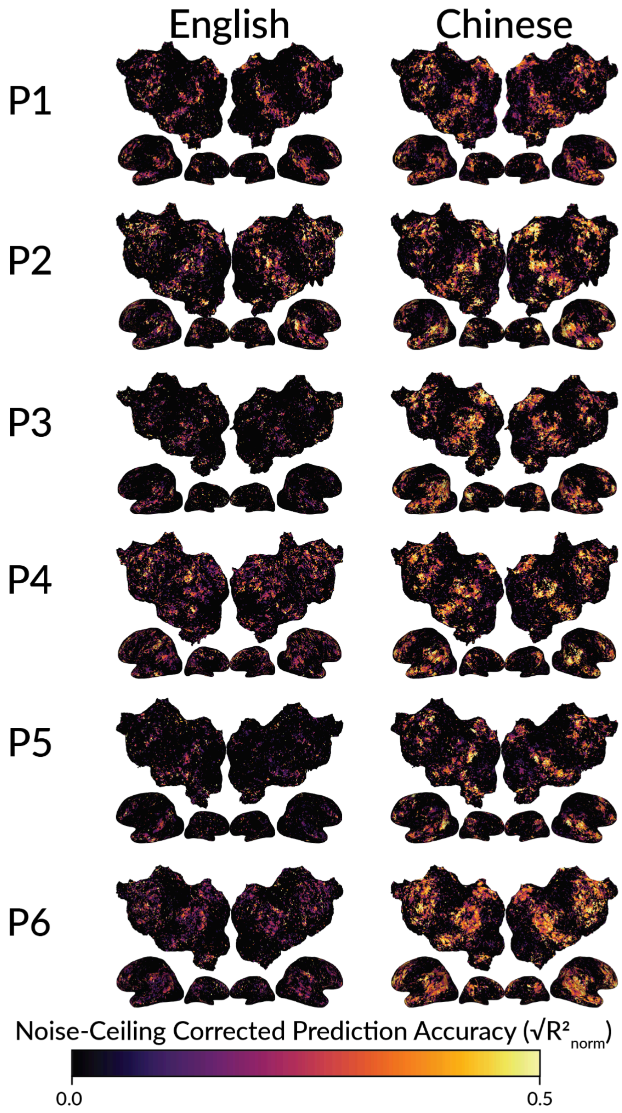


**Figure S8: Noise-ceiling corrected prediction accuracy for English vs Chinese.**
To determine whether differences in the noise-ceiling could explain the difference in prediction accuracy between languages, we computed the noise-ceiling corrected prediction accuracy ($R^2_{norm}$) for each language and participant separately. FastText was used as the semantic feature space. Noise-ceiling corrected prediction accuracy is shown on the flattened cortical surface of each participant's native brain space. The color of each voxel indicates prediction accuracy according to the colorbar at the bottom. For each participant, noise-ceiling corrected prediction accuracy is significantly higher in Chinese than in English (one-sided p<.05 by a permutation test). This suggests that the semantic model explains a higher proportion of total explainable signal in brain responses in Chinese than in English.

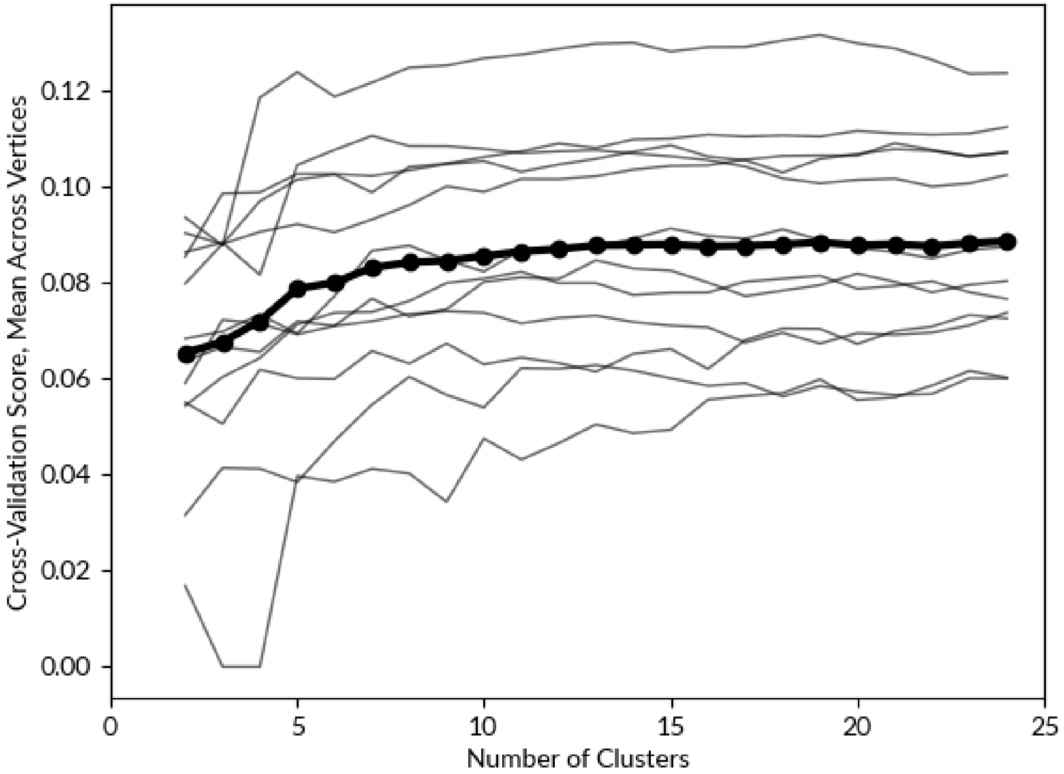


**Figure S9: Choice of number of clusters to use for semantic clustering**. Model weights were estimated for the semantic feature space, separately for each language and participant. A leave-one-out cross-validation approach was used to determine the optimal number of clusters into which to cluster model weights. In total, there are 12 sets of model weights across the six participants and two languages. Each set of model weights was projected to the template space. For each cross-validation fold, one of the twelve sets of model weights was held out. For each vertex the remaining eleven sets of model weights were averaged across participants and languages, and hierarchical clustering was used to obtain $N$ group-level clusters, separately for $N$ ranging from 2 through 25. For each number $N$ of clusters, the group-level model weight clusters were used to predict BOLD responses for the held out participant and language in vertex space. The cross-validation accuracy was computed for each vertex as the CD $R^2$ between the predicted and recorded BOLD responses. Only vertices that were well-predicted in both languages (group-averaged $\sqrt{R^2_{en}} > 0.05$ and $\sqrt{R^2_{zh}} > 0.05$) were included in this analysis. The mean $\sqrt{R^2}$ over vertices is shown for each number $N$ of clusters. Each thin line indicates the cross-validation scores for one fold. The bolded line indicates the mean across folds. The cross-validation score plateaus around five clusters. Thus, we used five semantic clusters for the analyses in Figure 3 and Figure 5.

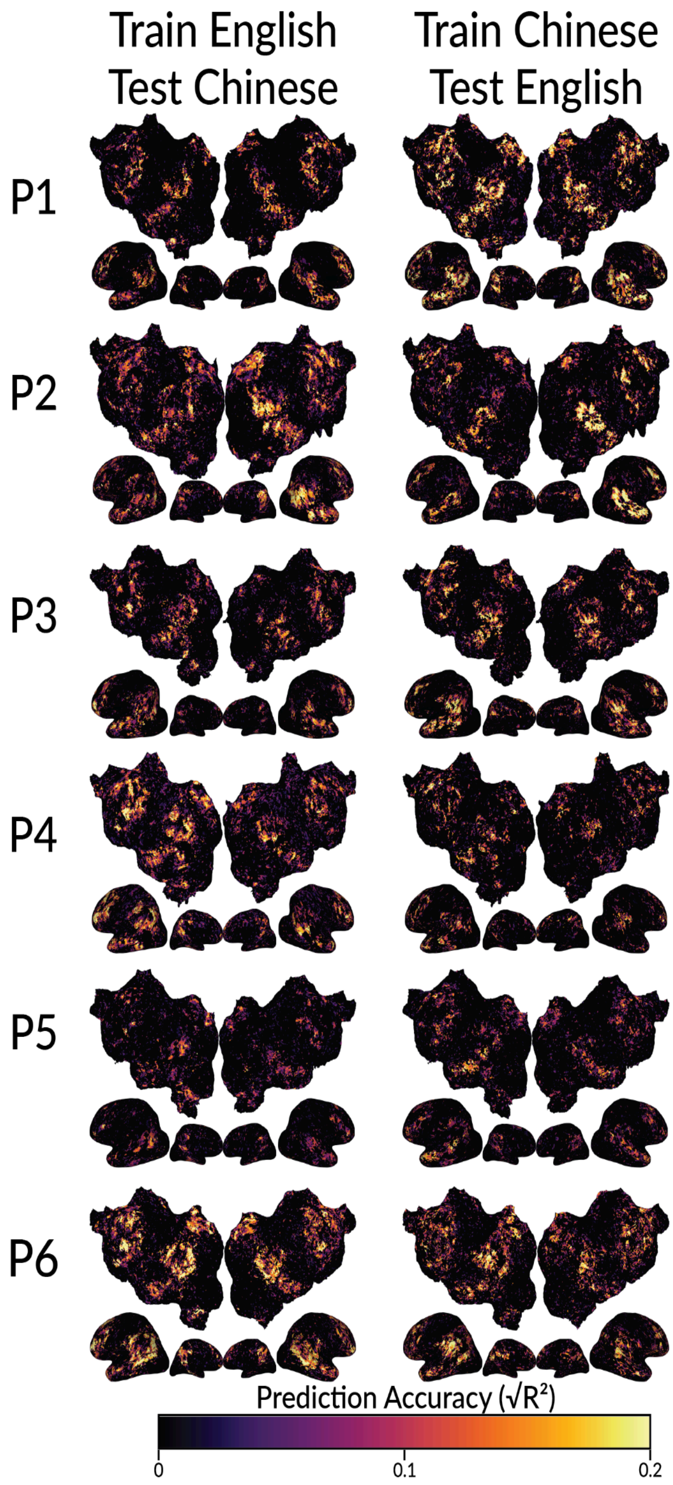


**Figure S10: Shared semantic representations between languages as shown by across-language prediction accuracy.** Across-language prediction accuracy is shown for each participant on the flattened cortical surface of the participant's native brain space. FastText was used as the semantic feature space. Estimated voxelwise model weights in one language were used to predict the held-out dataset in the other language. FastText was used as the semantic feature space. Prediction accuracy was computed as the CD ($R^2$) between predicted and recorded BOLD responses. Prediction accuracy is given by the color scale. Well-predicted voxels appear brighter. In each participant, the semantic model predicted in one language accurately predicts voxel responses to the other language throughout the semantic system. Thus, semantic representations within the semantic system are largely shared between languages.

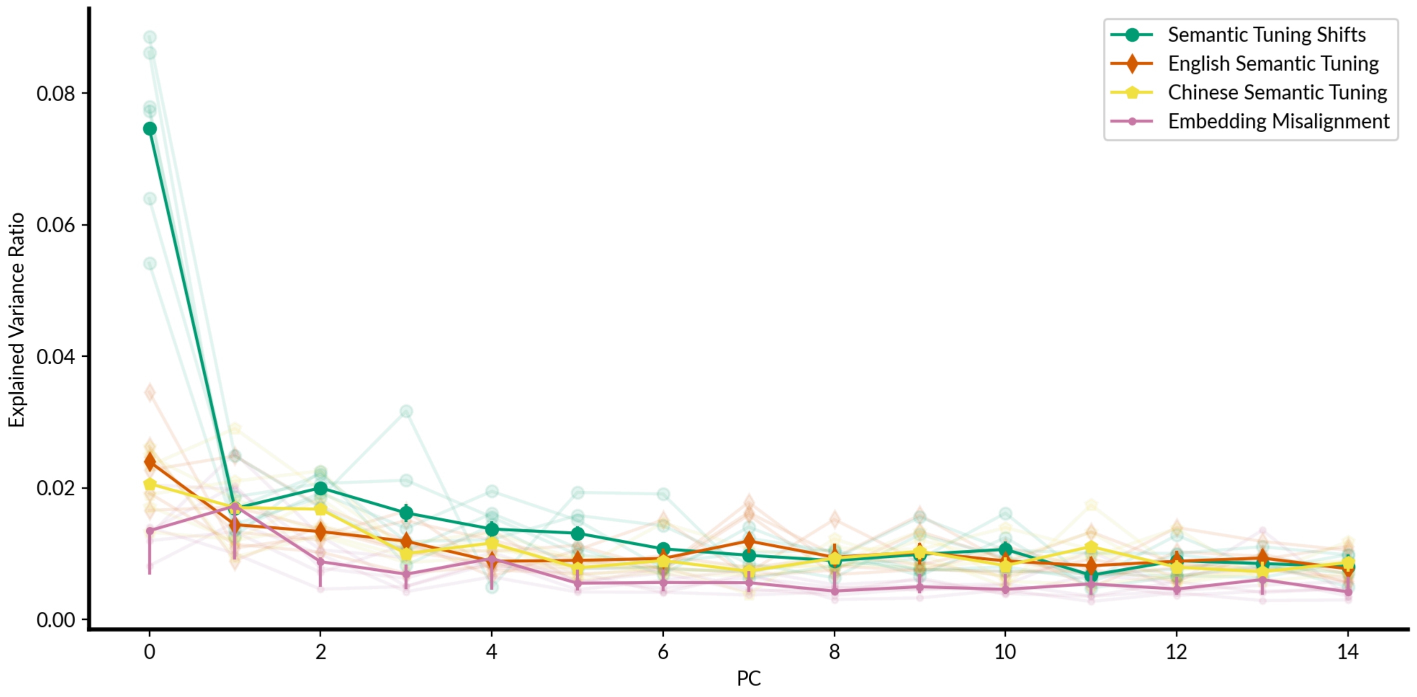


**Figure S11: Amount of variance explained by semantic tuning shift principal components (PCs).** Principal component analysis (PCA) was used to obtain the major dimensions of variation in semantic tuning shifts. To determine how many dimensions reliably capture variation in semantic tuning shifts a leave-one-participant-out procedure was used. At each step, one of the participants was left out and the voxelwise semantic tuning shifts from the other five participants were used to compute the principal components (PCs) of semantic tuning shifts (partial-group semantic tuning shift PCs). The partial-group semantic tuning shift PCs were used to compute the ratio of variance explained in the semantic tuning shifts of the left-out participant (green lines). Semantic tuning shift PCs were compared to three other PCs. First, the English semantic tuning PCs were computed by concatenating across voxels the 300-dimensional vectors of estimated model weights in English and then applying PCA across voxels to the resulting matrix. The English semantic tuning PCs were used to compute the ratio of variance explained in the semantic tuning shifts of each participant (orange lines). Second, the Chinese semantic tuning PCs were computed similarly to the English semantic tuning PCs but using the estimated semantic models weights in Chinese. The English semantic tuning PCs were used to compute the ratio of variance explained in the semantic tuning shifts of each participant (yellow lines). Third, the embedding misalignment PCs were computed by subtracting the embedding of each English word from its Chinese counterpart and then concatenating the 300-dimensional difference vectors across word pairs. Then PCA was applied across voxels to the embedding misalignment matrix. The embedding misalignment PCs were used to compute the ratio of variance explained in the semantic tuning shifts of each participant (pink lines). Transparent lines show variance explained in semantic tuning shifts for each individual participant, and opaque lines show the mean explained variance ratio across all participants. A bootstrapping approach was used to obtain confidence intervals for the variance explained by each PC. The voxel population was resampled 1000 times with replacement, and the tuning shift PCs were recomputed for each bootstrap iteration. 95% confidence intervals are shown by the error bars. The embedding misalignment PCs and the semantic tuning PCs for each language all explain significantly less variance than the first semantic tuning shift PC. This suggests that the first semantic tuning shift PC reliably captures variation in semantic tuning shifts, and that this dimension of tuning shifts is not merely an artifact of misalignments between embeddings for different languages, or of the primary dimensions of semantic tuning within each language.

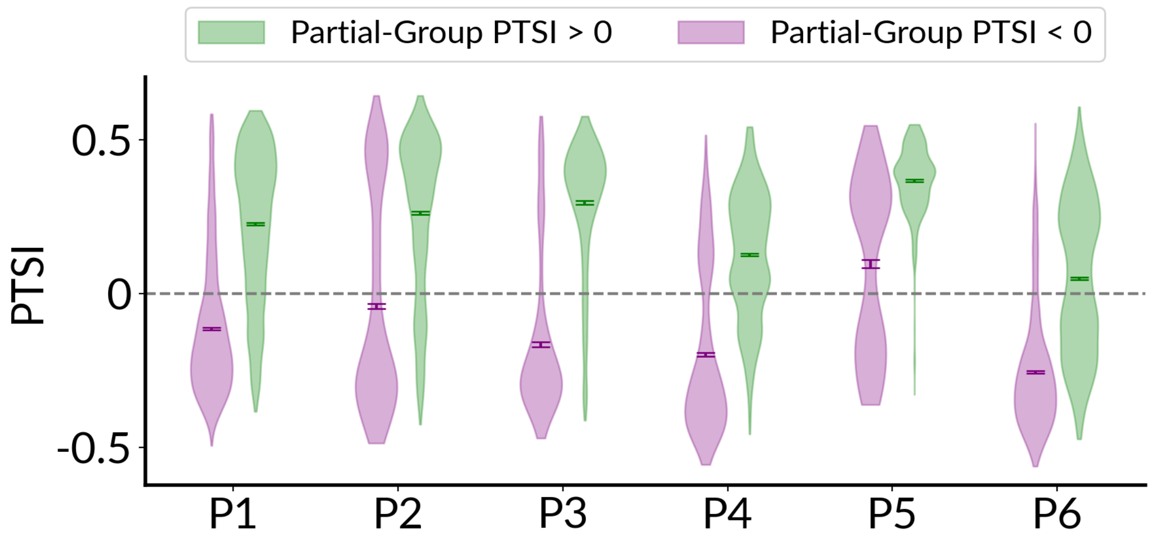


**Figure S12. Consistency of PTSI between participants.** Voxelwise PTSI of each participant was projected onto the template space. Then we hold out each participant, and use the other five participants to compute a group-level estimate of PTSI for each vertex. For each participant, violinplots show the distribution of vertexwise PTSI, separately for vertices in which PTSI is negative (purple violinplots) and positive (green violinplots) in the rest of the group. Vertices that were not well-predicted in both languages ($\sqrt{R^2} < 0.1$) were excluded from this analysis. Vertices with positive PTSI in the rest of the group also have positive PTSI in the participant ($p<.05$ by a one-sided t-test after Fisher z-transformation). Vertices with negative PTSI in the rest of the group also have negative PTSI in the participant ($p<.05$ by a one-sided t-test after Fisher z-transformation; except P5). Overall, these results show that the cortical distribution of PTSI is consistent across participants.

\- Primary Tuning Shift Dimension +

| - | + |
|---|---|
| 既 - already | 路边 - roadside |
| 之 - of | 山沟 - ravine |
| 位 - position | 砾石 - gravel |
| 以 - by | 不速之客 - uninvited guest |
| 更 - more | 打电话 - make a phone call |
| 是 - yes | 山顶 - hilltop |
| 起 - rise | 忍不住 - can't help it |
| 全 - all | 阿拉巴马州 - alabama |
| 打 - hit | 印第安人 - Indian person |
| 名 - name | 德克萨斯 - texas |
| 份 - portion | 研究生 - postgraduate |
| 令 - command | 所作所为 - behavior |
| 但 - however | 基督教 - Christianity |
| 七 - seven | 亚特兰大 - Atlanta |
| 和 - together | 有生以来 - for one's whole life |

**Figure S13: Interpretation of the first semantic tuning shift PC based on Chinese stimulus words.** Chinese stimulus words that best match each end of the PC are shown. Words closest to the negative end of the PC are shown in purple. Words closest to the positive end of the PC are shown in green. English translations are listed next to each word. The Chinese words closest to the negative end of the dimension are generally related to numbers. The Chinese words closest to the positive end of the PC are generally related to actions, people, and places. Interpretation of the PC is broadly consistent whether interpretations are based on Chinese stimulus words (shown here) or English stimulus words (shown in Figure 4A).

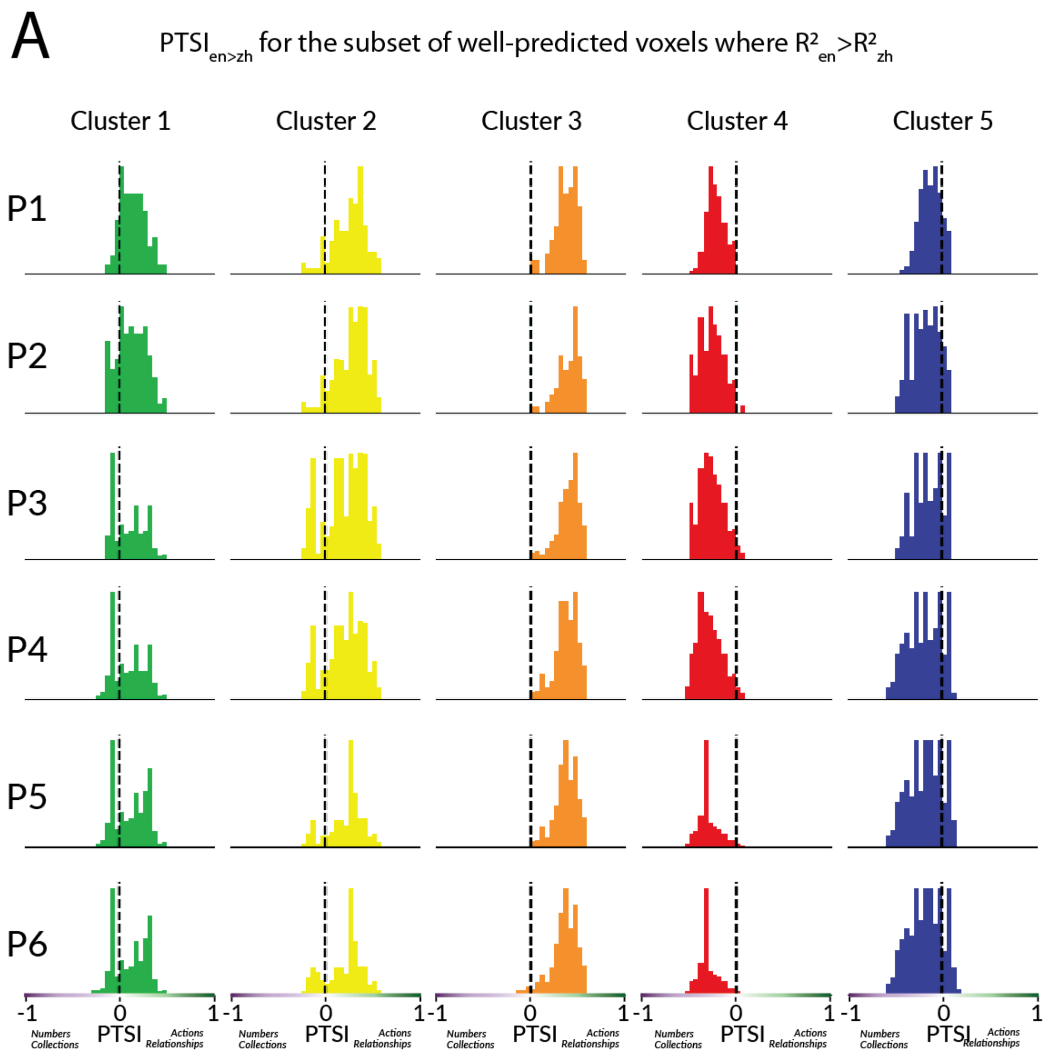

A
PTSI en>zh for the subset of well-predicted voxels where R² en > R² zh
Cluster 1
Cluster 2
Cluster 3
Cluster 4
Cluster 5
P1
P2
P3
P4
P5
P6
-1
0
1
PTSI
Numbers Collections
Actions Relationships

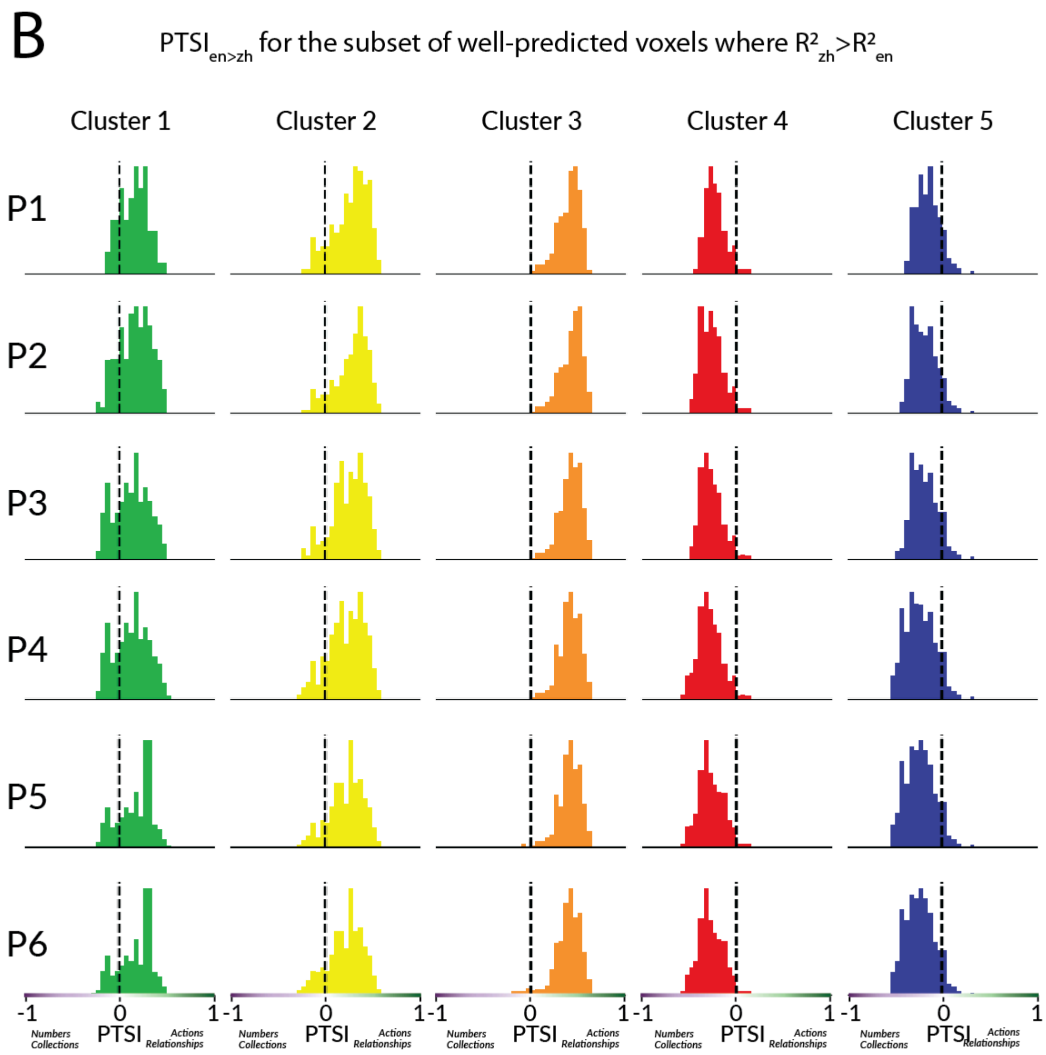
B
PTSI en>zh for the subset of well-predicted voxels where R² zh>R² en
Cluster 1
Cluster 2
Cluster 3
Cluster 4
Cluster 5
P1
P2
P3
P4
P5
P6
-1
0
1
PTSI
Numbers Collections
Actions Relationships

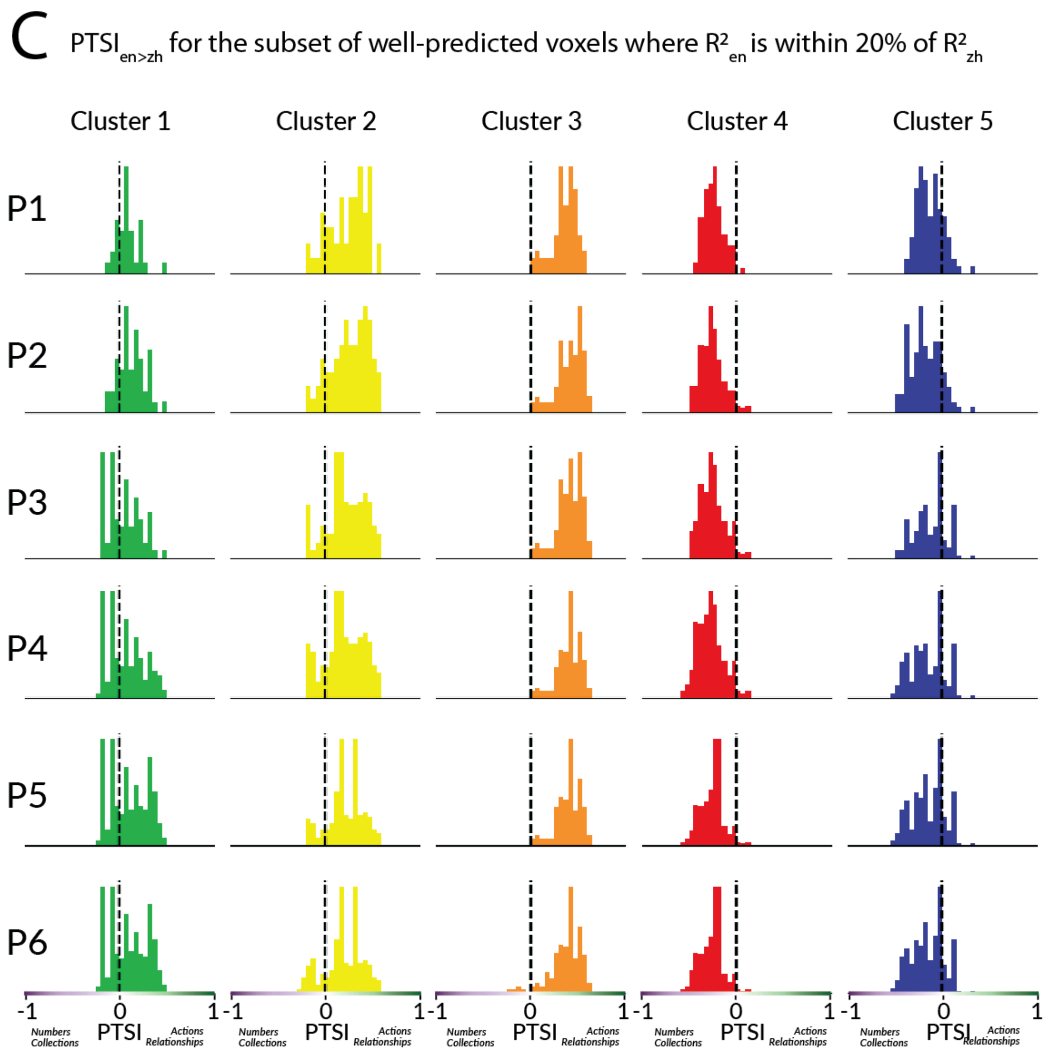


**Figure S14. Semantic tuning shifts for different voxel subsets.** To test whether the observed semantic tuning shifts are an artifact resulting from prediction accuracies being higher in Chinese than in English, we re-computed the first semantic tuning shift PC and voxelwise PTSI using three different subsets of well-predicted voxels: the subset where prediction accuracy is higher in English than in Chinese ($R^2_{en} > R^2_{zh}$; we refer to the resulting PC and PTSI as $PC_{en>zh}$ and $PTSI_{en>zh}$), the subset where prediction accuracy is higher in Chinese than in English ($R^2_{zh} > R^2_{en}$; we refer to the resulting PC and PTSI as $PC_{zh>en}$ and $PTSI_{zh>en}$), and the subset where prediction accuracy is similar between the two languages ($|R^2_{en} - R^2_{zh}| < 0.2 \times R^2_{zh}$; we refer to the resulting PC and PTSI as $PC_{en\sim zh}$ and $PTSI_{en\sim zh}$). We refer to the PC and PTSI shown in the main paper as $PC_{all}$ and $PTSI_{all}$.

First, we tested whether the direction of the first semantic tuning shift PC is consistent between the three subsets of voxels. The correlation between $PC_{en>zh}$ and $PC_{zh>en}$ is 0.91 (p<.001). The correlation between $PC_{en\sim zh}$ and $PC_{all}$ is 0.98 (p<.001 by a one-sided permutation test). Furthermore, for each participant, the correlation between $PTSI_{en>zh}$ and $PTSI_{all}$, the correlation between $PTSI_{en>zh}$ and $PTSI_{zh>en}$, and the correlation between $PTSI_{en\sim zh}$ and $PTSI_{all}$ are at least 0.98. Thus, the direction of the first semantic tuning shift PC is not confounded by differences in model accuracy.

Next, we tested whether the distribution of PTSI across semantic clusters (Figure 5) is an artifact of differences in model accuracy. The panels in this figure show the distribution of voxelwise semantic tuning shifts for each semantic clusters. The format is similar to Figure 5 in the main paper. **A.** Histograms show the distribution of voxelwise $PTSI_{en>zh}$ for each cluster and participant separately. Voxels in Clusters 1, 2, and 3 have positive $PTSI_{en>zh}$ ($p<.05$ for each cluster by a two-sided t-test after Fisher z-transformation, for all participants except P3 Cluster 1 where p=.4, P6 Cluster 1 where p=0.4, and P6 Cluster 2 where p=.1). Voxels in Clusters 4 and 5 have negative $PTSI_{en>zh}$ ($p<.05$ for each cluster by a two-sided t-test after Fisher z-transformation, for all participants except P3 Cluster 5 where p=.6). The direction of these distributions is consistent with those shown in Figure 5, which included all well-predicted voxels. **B**. Histograms show the distribution of voxelwise $PTSI_{zh>en}$ for each cluster and participant separately. Voxels in Clusters 1, 2, and 3 have positive $PTSI_{zh>en}$ ($p<.05$ for each cluster by a two-sided t-test after Fisher z-transformation, for all participants). Voxels in Clusters 4 and 5 have negative $PTSI_{zh>en}$ ($p<.05$ for each cluster by a two-sided t-test after Fisher z-transformation, for all participants). The direction of these distributions is consistent with those shown in (A), and with those shown in Figure 5. **C**. Histograms show the distribution of voxelwise PTSI for each cluster and participant separately. For voxels in Clusters 1, 2, and 3 $PTSI_{en\sim zh}$ is positive ($p<.05$ for each cluster by a two-sided t-test after Fisher z-transformation; except Cluster 1 for P3 where p=.4 and voxels have negative $PTSI_{en\sim zh}$, Cluster 1 for P6 where p=.1, and Cluster 2 for P6 where p=.5). For voxels in Clusters 4 and 5 $PTSI_{en\sim zh}$ is negative ($p<.05$ for each cluster by a two-sided t-test after Fisher z-transformation; except Cluster 4 for P5 where p=.08, and Cluster 5 for P3 where p=.7). The direction of these distributions is consistent with those shown in (A), and with those shown in Figure 5 in the main paper. Overall, these results suggest that semantic tuning shifts shown in the main results are not confounded by differences in prediction accuracy.

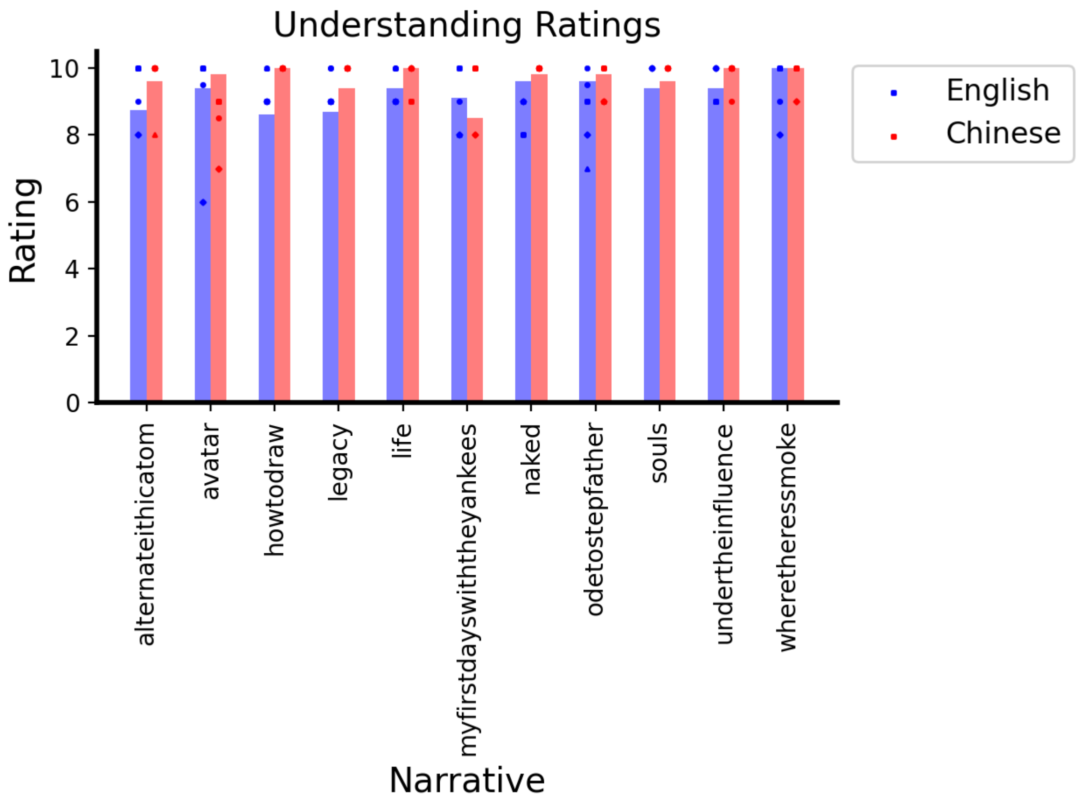


**Figure S15. Self-reported comprehension scores.** After each narrative, participants were asked to report how well they understood the narrative on a scale from 1 (lowest) to 10 (highest). Participants' scores are shown for English (blue) and Chinese (red) narratives separately. Each dot reflects one participant's score. Each bar reflects the mean across participants. Participants' ratings were high for both languages (97% of ratings were 8 or above), suggesting that participants understood the stories well in both languages. Ratings were slightly higher for Chinese than for English (p=.001 by a two-sided paired t-test). Ratings were not collected for participant P1 because of an update in scanning procedure. Among participants P2-P6, four ratings ( P2-"souls"-English, P4-"souls"-English, P6-"my first day with the yankees"-English, and P4-"souls"-Chinese) are missing due to a lack of time at the end of the scan.

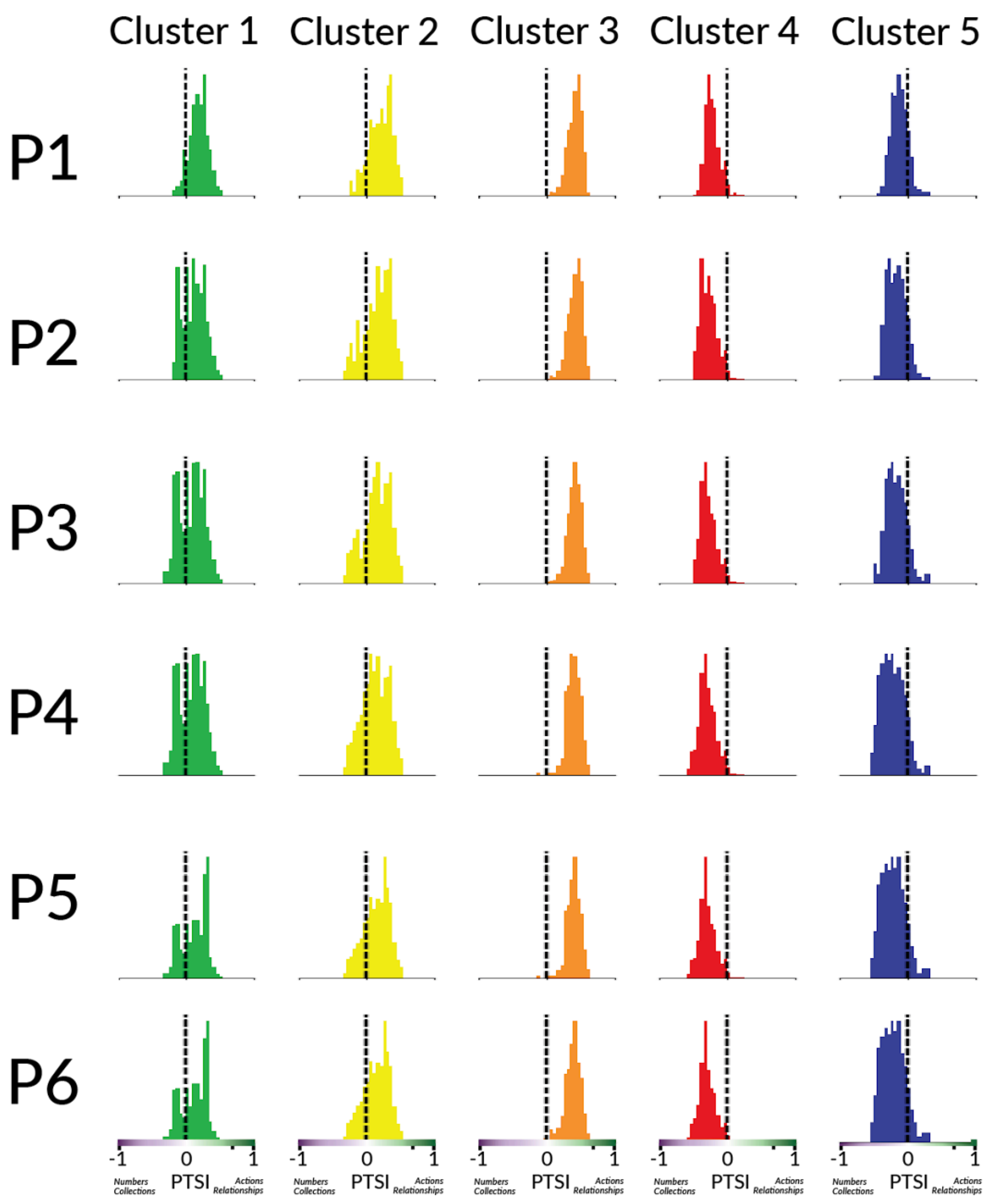


**Figure S16: Semantic tuning shifts for different clusters when Chinese model weights are used.** The semantic tuning shifts for different clusters depicted in Figure 5A were computed using English model weights. To test whether these results are independent of the language of model weights used for clustering, Chinese model weights are used to assign each voxel to a semantic cluster. Histograms indicate the distribution of PTSI values for voxels in each cluster, for each participant separately. Semantic tuning shifts for voxels in Clusters 1, 2, and 3 (green, yellow, and orange histograms; except P3 cluster 1 where $p=.52$) are positively correlated with the first semantic tuning shift PC ($p<.05$ for each cluster by a two-sided t-test after Fisher z-transformation). Semantic tuning shifts for voxels in Clusters 4 and 5 (red and blue histograms) are negatively correlated with the PC ($p<.05$ for each cluster by a two-sided t-test after Fisher z-transformation). These results show that the results shown in Figure 5A are consistent whether English or Chinese model weights are used to assign each voxel to a semantic cluster.

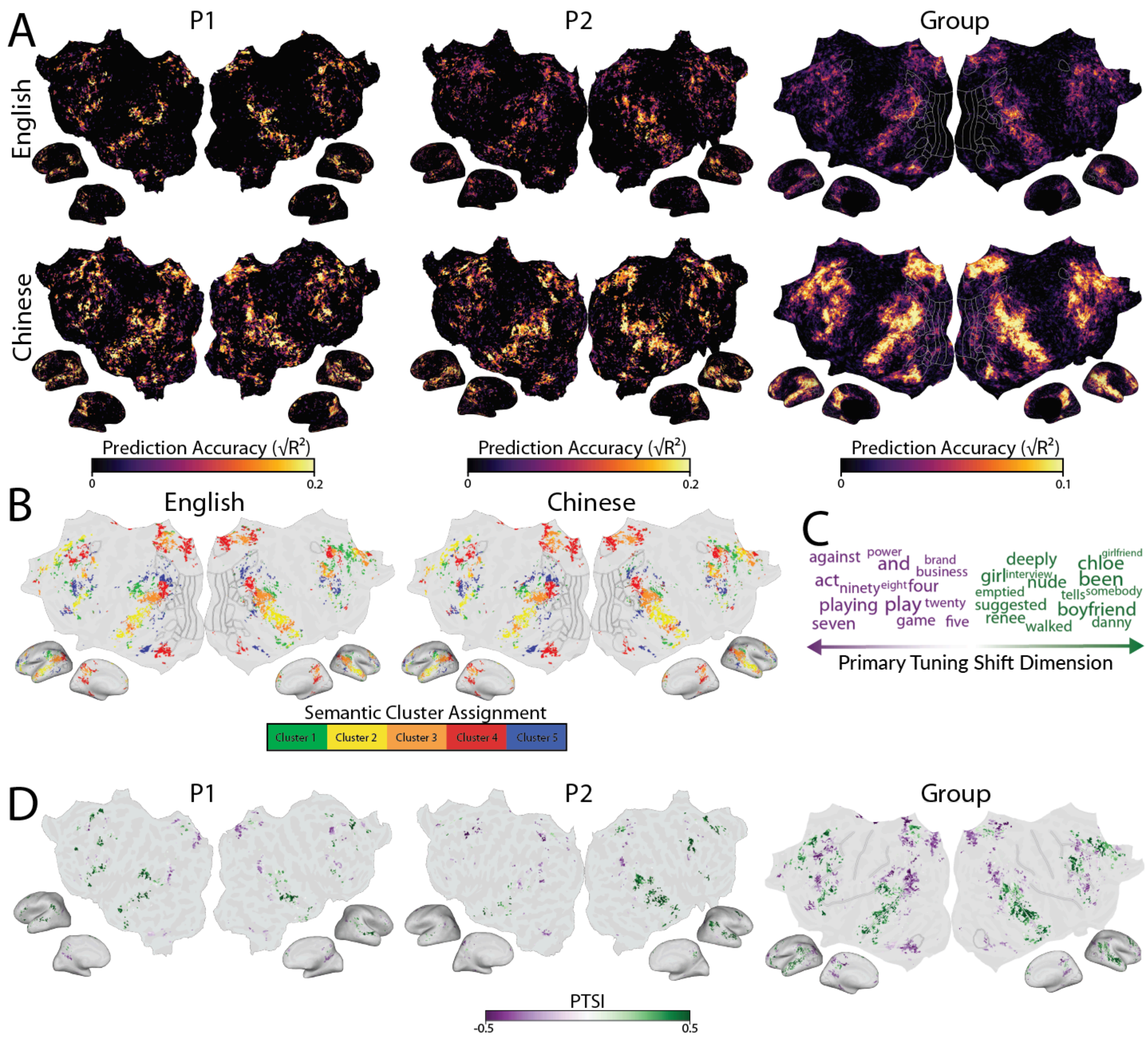


**Figure S17. Estimated semantic representations after additionally regressing out grammatical category and word/lemma frequency.** A. Prediction accuracy is shown for two representative participants (P1 and P2), and at the group level. Results are shown separately for each participant. For each of the six participants, prediction accuracy is highest in the bilateral temporal, parietal, and prefrontal cortices. Semantic information is represented in similar regions for both languages. B. Semantic cluster assignments. Results are shown at the group-level. Group-level cluster assignments are the same across languages in 80% of well-predicted vertices. For each individual participant, cluster assignments are the same across languages in over 70% of well-predicted voxels (P1 82%, P2 71%, P3 84%, P4 74%, P5 89%, P6 86%). Semantic cluster assignments are highly similar between languages C. Interpretation of the first semantic tuning shift PC. D. Distribution of PTSI is shown for two representative participants (P1 and P2), and at the group level. The results shown here are consistent with results shown in Figures 1-5. This suggests that the results are likely not confounded by word frequencies or grammatical categories.

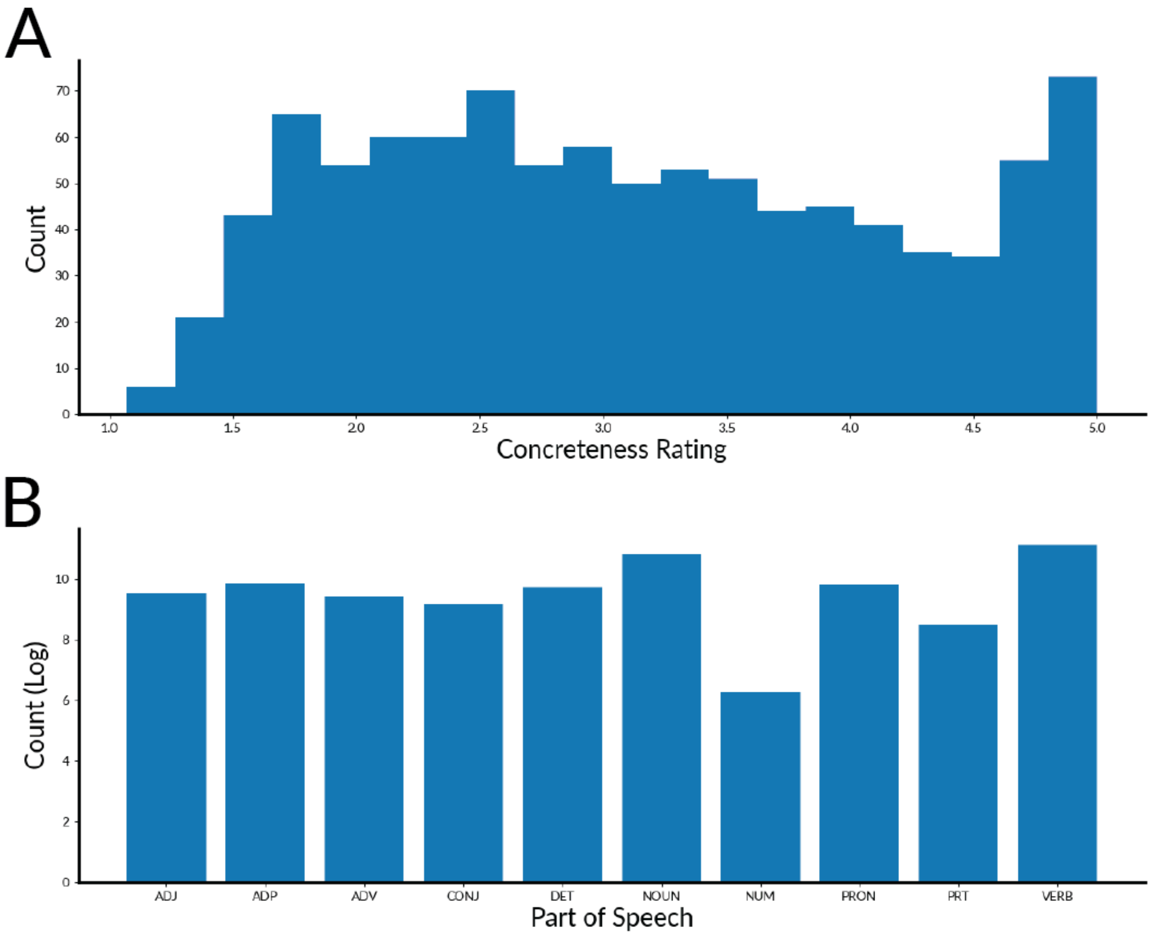


**Figure S18. Distribution of word concreteness and grammatical categories.** A. To estimate the distribution of word concreteness in the stimulus text, Brysbaert human ratings of word concreteness were used[82]. The histogram shows counts of word concreteness over all unique words in the original English narratives (1=very abstract, 5=very concrete; note that very few words were rated as very abstract (1.0-1.5) in the Brysbaert dataset). B. To estimate the distribution of grammatical categories in the stimulus text, the Berkeley neural parser was used to determine the part of speech of each word in the English narratives[83]. The histogram shows the logarithmic counts for each part of speech category. The stimulus narratives contain both concrete and abstract words, and a range of grammatical categories.

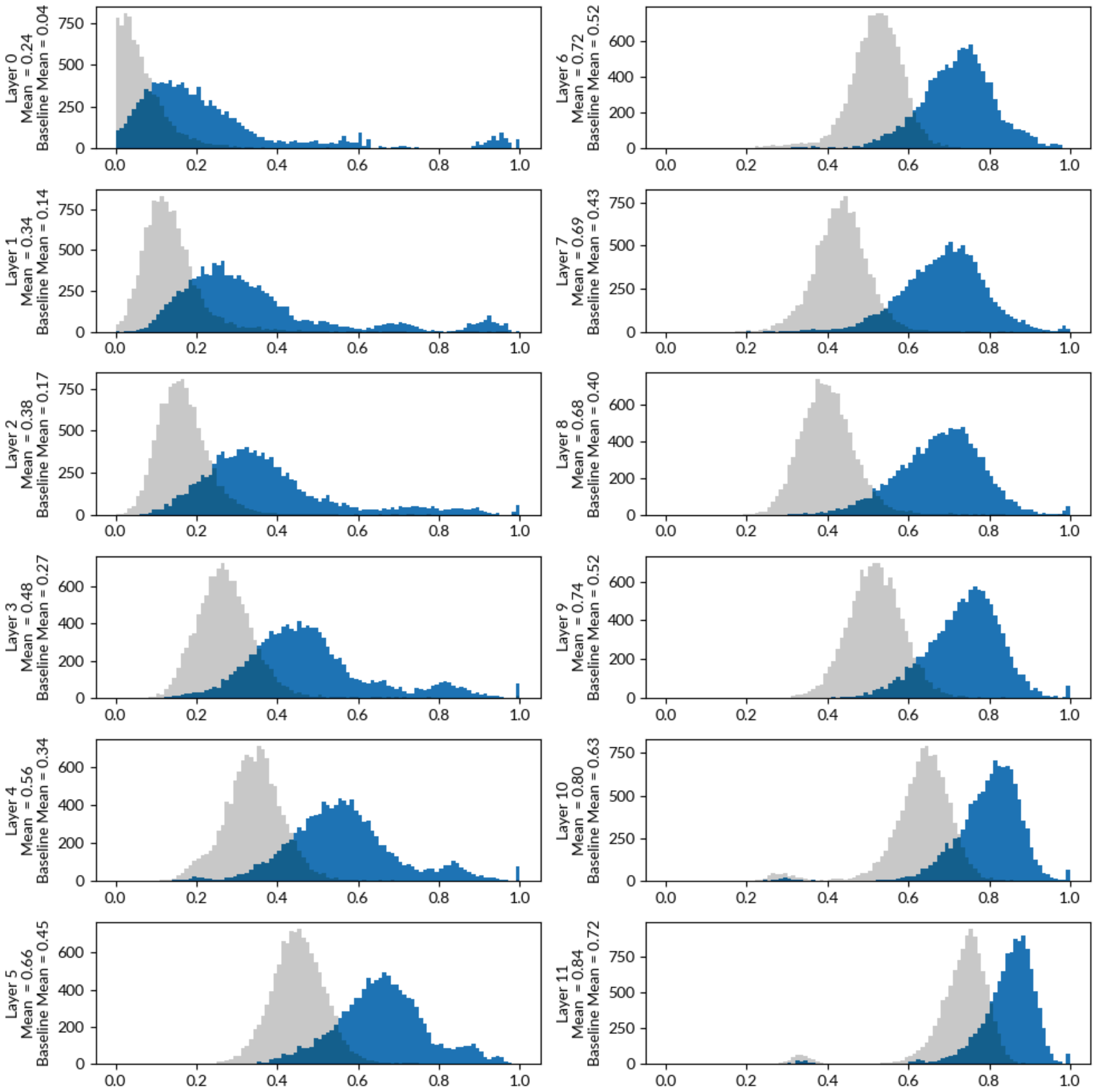


**Figure S19: Best-aligned layers of multilingual BERT (mBERT)**. To determine which layer of mBERT produces the best multilingual semantic embedding space, the alignment of each layer of mBERT was measured between English and Chinese. To evaluate alignment between mBERT embeddings of English and Chinese we used the TsinghuaAligner test dataset[97]. This dataset consists of 450 sentences that are provided in both English and Chinese, as well as manually annotated pairs that indicate pairs of English and Chinese words that mean the same thing. For each pair of words, each layer of mBERT was used to obtain two 768-dimensional embeddings: one for the English word, and one for the aligned Chinese word. For each layer separately, the similarity between the respective English and Chinese embeddings was measured as the cosine similarity between the two embeddings. As a baseline comparison, we measured the cosine similarity between the embeddings of randomly selected English-Chinese word pairs that did not mean the same thing. For each layer of mBERT we show the distribution of cosine similarities between aligned English-Chinese word pairs (blue) and the distribution of cosine similarities between randomly selected English-Chinese word pairs (grey). Embeddings for matched pairs of words become more similar in later layers. However, the latest layers produce outliers of poorly aligned word pairs that may bias semantic tuning shift estimates. Layer 9 produces the best aligned embeddings: embeddings of paired words have high cosine similarity relative to embeddings of random word pairs, without substantial outliers. Thus, we use layer 9 embeddings to validate the results from the fastText semantic embedding space (Supplementary Figures S1-S4).

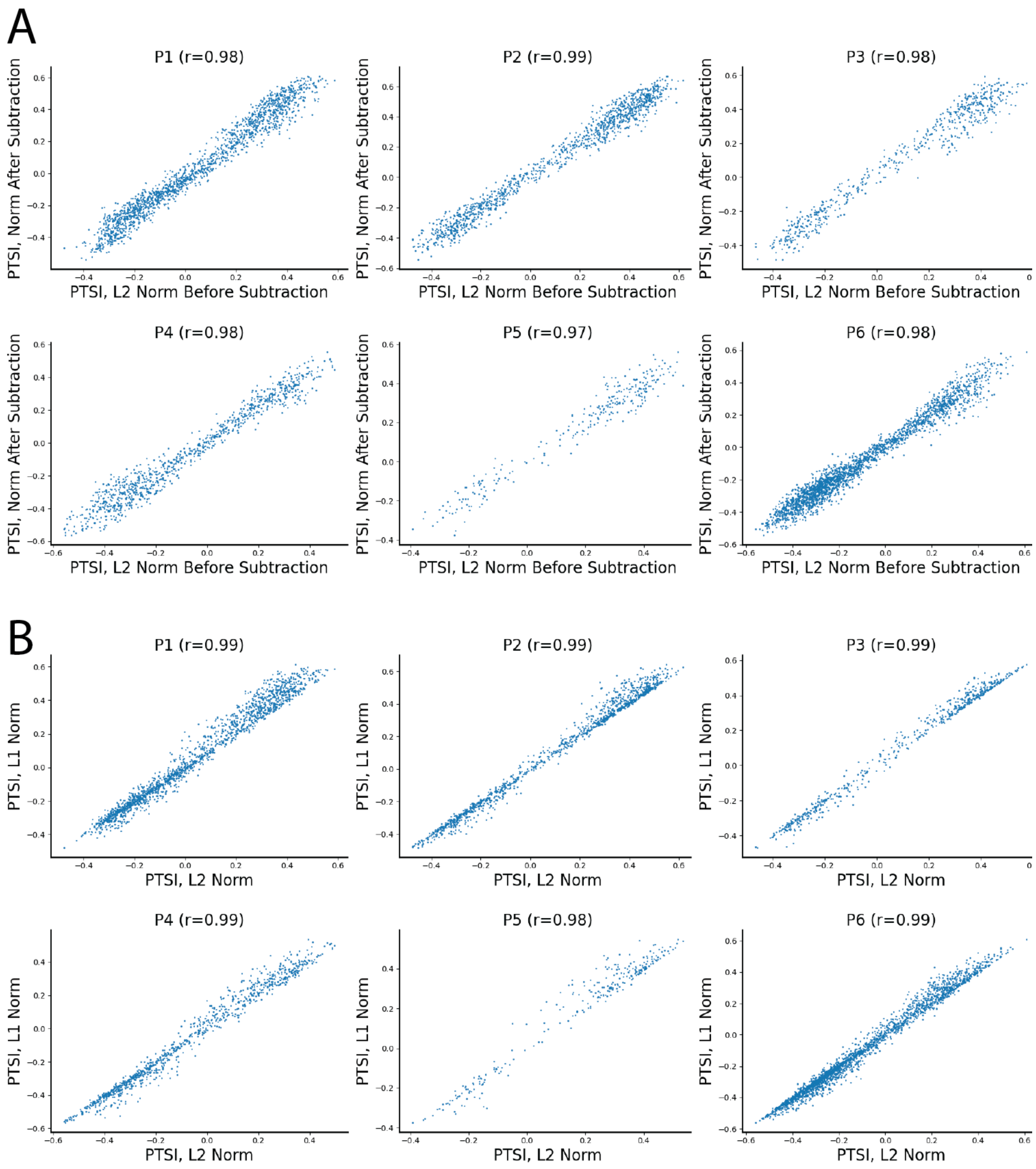


**Figure S20**. Comparison between PTSI normalization methods. A. PTSI comparison, L2 normalization before subtraction (x-axis) vs. L2 normalization after subtraction (y-axis). Each point reflects one voxel. Results are shown for each individual participant. The new and original PTSI values are highly correlated (r>.97 for all six participants). B. PTSI comparison, L2 normalization (x-axis) vs. L1 normalization (y-axis). The format is the same as in A. The new and original PTSI values are highly correlated (r>.97 for all six participants). These results suggest that the PTSI computation is not influenced by the specific normalization procedure.

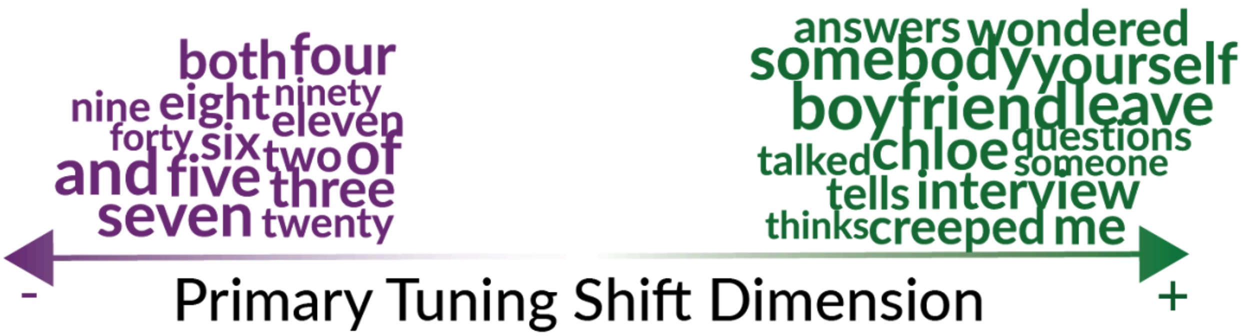


**Figure S21: First semantic tuning shift PC, based on significantly well-predicted voxels.** In Figure 4 the first semantic tuning shift PC is estimated using voxels that are well-predicted in both languages ($\sqrt{R^2} > 0.1$ in both languages). To verify that the estimated PC is robust to the voxel selection method, we instead estimated the PC using voxels that were significantly well-predicted in both languages (one-sided p<.05, after Benjamini-Hochberg correction for multiple comparisons). The words that best match the significance-based estimate of the PC are shown. This dimension separates number/collection-related semantics (purple) from action/relationship-related semantics. The Pearson correlation between the significance-based estimate and the prediction accuracy-based estimate of the PC is 0.97. Thus, the estimate of the first semantic tuning shift PC is robust to the voxel selection method.

| Participant | P1 | P2 | P3 | P4 | P5 | P6 |
|---|---|---|---|---|---|---|
| Age when you began ACQUIRING language 1 | 1 month | 2 | since born | Since born | 0 | Native |
| Age when you began ACQUIRING language 2 | 11 years old | 7 | 4 | 6 | 2 | 6 |
| Age when you became FLUENT in language 1 | 7 years old | 7 | 6 | 5 | 5 | Native |
| Age when you became FLUENT in language 2 | 24 years old | 18 | 20 | 18 | 16 | 19 |
| How many years and months have you spent in a COUNTRY where language 1 is spoken? (E.g. 2 years 3 months) | 18 years | 18 years | 18 years | 20 years | 18 years | 18 years |
| How many years and months you spent in a COUNTRY where language 2 is spoken? (E.g. 2 years 3 months) | 12 years | 6 years 1 months | 6 years and 9 months | 5 years | 6 years | 6 years 6 months |
| How many years and months have you spent in a FAMILY where language 1 is spoken? (E.g. 2 years 3 months) | 30 years | 18 years | 18 years | 25 years | 18 years | 18 years |
| How many years and months you spent in a FAMILY where language 2 is spoken? (E.g. 2 years 3 months) | 8 years | 0 | 0 | 1 month | 0 | 0 |
| How many years and months have you spent in a SCHOOL and/or WORKING environment where language 1 is spoken? (E.g. 2 years 3 months) | 18 years | 11 years | 12 years | 20 years | 12 years | 18 years |
| How many years and months you spent in a SCHOOL and/or WORKING environment where language 2 is spoken? (E.g. 2 years 3 months) | 12 years | 6 years 1 months | 6 years and 9 months | 5 years | 6 years | 6 years 6 months |
| Please circle to what extent you are CURRENTLY EXPOSED to language 1 in INTERACTING WITH FRIENDS: | 5 | 4 | 7 | 7 | 5 | 7 |
| Please circle to what extent you are | 10 | 4 | 5 | 7 | 7 | 5 |

| | | | | | | |
|---|---|---|---|---|---|---|
| CURRENTLY EXPOSED to language 2 in INTERACTING WITH FRIENDS: | | | | | | |
| Please circle to what extent you are CURRENTLY EXPOSED to language 1 in INTERACTING WITH FAMILY: | 8 | 10 | 3 | 10 | 3 | 10 |
| Please circle to what extent you are CURRENTLY EXPOSED to language 2 in INTERACTING WITH FAMILY: | 6 | 0 | 0 | 0 | 0 | 0 |
| Please circle to what extent you are CURRENTLY EXPOSED to language 1 at SCHOOL | 0 | 2 | 0 | 0 | 0 | 0 |
| Please circle to what extent you are CURRENTLY EXPOSED to language 2 at SCHOOL | 10 | 8 | 10 | 10 | 10 | 9 |
| Please circle to what extent you are CURRENTLY EXPOSED to language 1 at WORK | 1 | 0 | 0 | 0 | 0 | 2 |
| Please circle to what extent you are CURRENTLY EXPOSED to language 2 at WORK | 10 | 8 | 10 | 10 | 10 | 9 |

**Table S1.** Participant responses to language use questionnaire.
